# 3D-bioprinted marine bacteria for the degradation of bioplastics

**DOI:** 10.1101/2025.02.05.636490

**Authors:** Luying He, Hongyi Cai, Ram S. Gona, Manasi S. Gangan, Timothy Lai, Meredith N. Silberstein, Anne S. Meyer

## Abstract

The severe, long-lasting harm caused by plastic pollution to marine ecosystems and coastal economies has led to the development of biodegradable plastics; however, their limited decomposition in cold, dark marine environments remains a challenge. Here, we present our newly developed technologies for creating 3D-bioprinted living materials for bioplastic degradation with specific use in marine environments. Our approach integrates halotolerant bioplastic-degrading bacterium *Bacillus* sp. NRRL B-14911 into alginate-based bio-ink to print an engineered living material (ELM) termed a “bio-sticker.” Quantification of bacteria viability reveals that bioprinted marine bacteria survive within bio-stickers for more than three weeks. The rate at which the bio-stickers degrade the bioplastic polyhydroxybutyrate (PHB) can be tuned by altering bio-sticker biomass concentration, bioplastic concentration, or incubation temperature. Bio-stickers that are transferred to a new PHB sample still retain high biodegradation activity, demonstrating their durability. Strain sweep oscillatory tests demonstrate viscoelastic behavior of the bio-stickers. Monotonic tensile tests indicate that the elastic modulus and the adhesion of the bio-stickers are not negatively impacted by bacteria growth or incubation temperature. Our work paves the way for development of ELMs to facilitate the inclusion of bioplastics within the blue economy, promoting the emergence of more sustainable and eco-friendly materials.

## 1. Introduction

The rapid escalation of marine plastic pollution presents a critical threat to both marine ecosystems and coastal economies. Plastic waste has been discovered in nearly all marine ecosystems, ranging from surface waters to the deepest trenches.^[1]^ An estimated 5-13 million metric tons of land-based plastic waste enter the ocean on a yearly basis, an amount predicted to increase sharply in upcoming years.^[2]^ Current projections estimate that the oceans will contain more plastic than fish by weight by 2050, if the rate of plastic buildup in the ocean continues unabated.^[3]^ Bioplastics, plastics formed from biologically grown polymers rather than fossil resources, have attracted attention as potential mitigating solutions to plastic waste and are a quickly growing sector of the plastic market. However, bioplastics currently comprise less than 1% of the plastics industry.^[4]^ While bioplastics such as polylactic acid (PLA) have been designed and tested for disposal in industrial composting facilities, these “biodegradable” bioplastics show limited biodegradation in cold, dark ocean environments.^[5]^ As a result, new sustainable materials with demonstrated biodegradability in marine conditions are urgently needed to preserve marine ecosystems.

Polyhydroxyalkanoates (PHA) are a highly promising class of bioplastics since they share similar mechanical properties to traditional rigid structural plastics such as polypropylene.^[6]^ PHAs are polyesters that are naturally synthesized and accumulated within bacteria cells in order to store energy and carbon resources,^[7]^ and their specific chemical structure and material properties can vary depending on the specific producer organism and environmental conditions. Poly-3-hydroxybutyrate (PHB) is one of the most common types of PHA and is produced by a variety of bacteria species.^[8]^ In contrast to traditional plastics and many types of bioplastics, PHB is fully biodegradable and can be broken down to H_2_O and CO_2_ without creating microplastic waste.^[8a]^ PHB biodegradation is accomplished by a variety of microbes that express PHB depolymerizing enzymes, and it can be degraded in both aerobic and anaerobic environments.^[9]^ Importantly, PHB is able to be biodegraded in saltwater environments,^[10]^ showing up to 88-99% biodegradation after 49 days in seawater.^[10a]^ The development of new technologies that allow for accelerated or tunable PHB degradation rates in marine environments could enable this bioplastic to be employed in a diverse range of different aquatic applications.

To facilitate the rapid degradation of bioplastics in marine environments, we have developed new engineered living materials (ELMs) designed for targeted and controlled degradation of plastic waste. Engineered living materials are a class of next-generation functional materials composed of engineered biological systems that create, modify, or maintain their own material structures or properties.^[11]^ Utilizing the transformative technology of 3D-bioprinting, we have designed three-dimensional engineered living material structures with PHB-depolymerization ability. Our custom-built 3D bioprinter can deposit a specialized “bio-ink” composed of alginate, which is a bio-compatible hydrogel matrix, and living microbes.^[12]^ Upon extrusion onto a surface containing chemical cross-linkers, the bio-ink undergoes alginate polymerization, resulting in a solid, three-dimensionally patterned hydrogel structure that contains embedded bacterial cells. We hypothesized that the incorporation of microbes derived from ocean ecosystems would create living materials that can retain their viability and metabolic activities in saltwater environments. A number of marine microorganisms have been identified that are able to produce exodepolymerase enzymes active against PHB plastics.^[13]^ Among these is *Bacillus* sp. NRRL B-14911,^[13d]^ a bacterium that secretes the PhaZ enzyme. PhaZ can break down long polymeric PHB chains into HB monomers, which the microbe can subsequently metabolize.^[14]^ Incorporation of *Bacillus* sp. NRRL B-14911 into 3D-bioprinted materials will allow for automated production of living “bio-stickers” that support spatially-distributed degradation of PHB plastics throughout an extended material lifetime.

In this work, we screened several strains of marine microbes to select a strain with viability and efficient PHB-depolymerization ability within polymerized bio-ink alginate matrices in seawater-mimicking media. Colony forming unit (CFU) assays indicated that *Bacillus* sp. NRRL B-14911 retained viability and PHB-degradation activity within a 3D-bioprinted alginate matrix for up to four weeks post-printing. Alteration of bio-printing parameters including biomass within the bio-ink, incubation temperature, and PHB concentration was shown to tune the rate of PHB degradation by the 3D-printed bio-stickers. The bio-stickers were able to be transferred to fresh PHB substrates after 1-2 weeks and retained similar PHB depolymerization activity to fresh bio-stickers. Analysis of the mechanical properties of the bio-stickers revealed viscoelastic behavior and showed that extended incubation of the bio-stickers did not significantly affect their elastic modulus or adhesion to PHB. Our 3D-bioprinted living bio-stickers address the challenge of biodegradable plastics in cold marine environments and will open new avenues for integrating eco-friendly, sustainable materials into the blue economy.

## 2. Results

PHB-degrading bio-stickers were created by patterning a bio-ink mixture of living microbes and alginate polymer in three dimensions onto a surface supplemented with calcium ions (Ca^2+^) (Figure 1). The Ca^2+^ ions promote cross-linking of the alginate chains, solidifying the bio-ink into a flexible, adhesive bio-sticker. The bio-sticker provides mechanical stability as well as a hydrated environment for growth and function of PHB-degrading marine microbes, allowing them to depolymerize PHB materials to which the bio-sticker is adhered over extended time periods.

**Figure 1.**
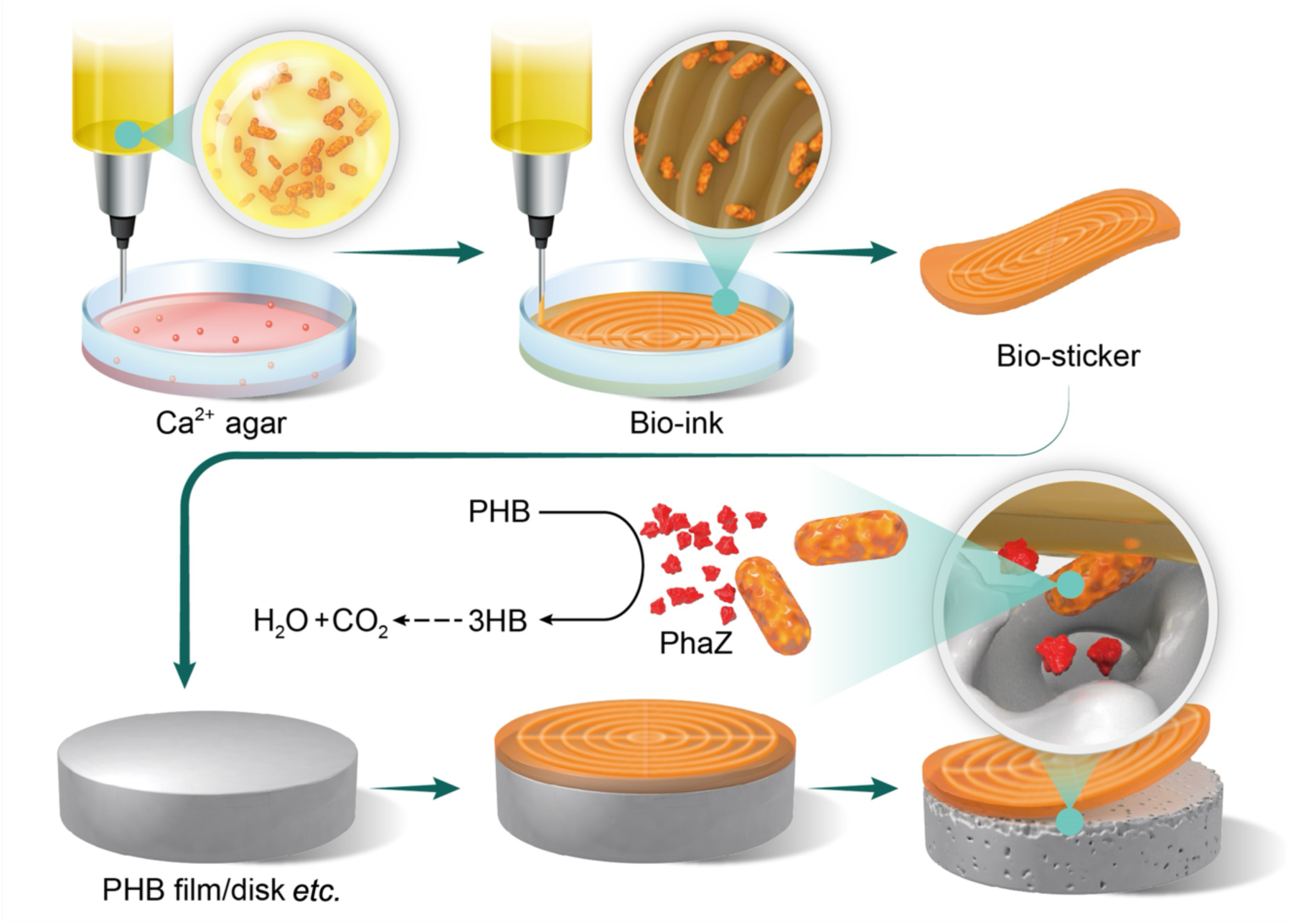
3D bioprinting of marine bacteria for biodegradation of bioplastic. Bio-ink consisting of marine bacteria and alginate is patterned in 3D onto an agar surface containing Ca^2+^. Interaction of bio-ink with Ca^2+^ ions drives the cross-linking of alginate chains to form a solid bio-sticker that can be transferred onto a bioplastic sample. The embedded microbes produce and excrete PhaZ enzyme to catalyze biodegradation of PHB bioplastic into HB monomers, which are further metabolically decomposed into H_2_O and CO_2_. The bottom panel shows biodegradation of PHB powder by patterned bio-stickers over a 12-day period, as seen by the formation of an expanding clear zone.

### 2.1 Design of 3D-bioprinted bio-stickers with PHB-degradation activity

To create 3D-printed engineered living materials with PHB degradation activity in marine environments, bacteria strains were screened for their ability to degrade PHB in marine-like environments. Four bacteria strains were selected that were isolated from marine sources and had been observed to demonstrate PHB-depolymerization ability: *Bacillus* sp. NRRL B-14911,^[14–15]^ *Comamonas testosteroni*,^[16]^ *Marinobacter* sp. NK-1,^[17]^ and *Microbulbifer* sp. SOL66.^[18]^ Growth curves recorded over a 24-hour incubation in Marine Broth at 30°C indicated that all four strains were able to proliferate in ocean-mimicking conditions (Figure 2A).

**Figure 2.**
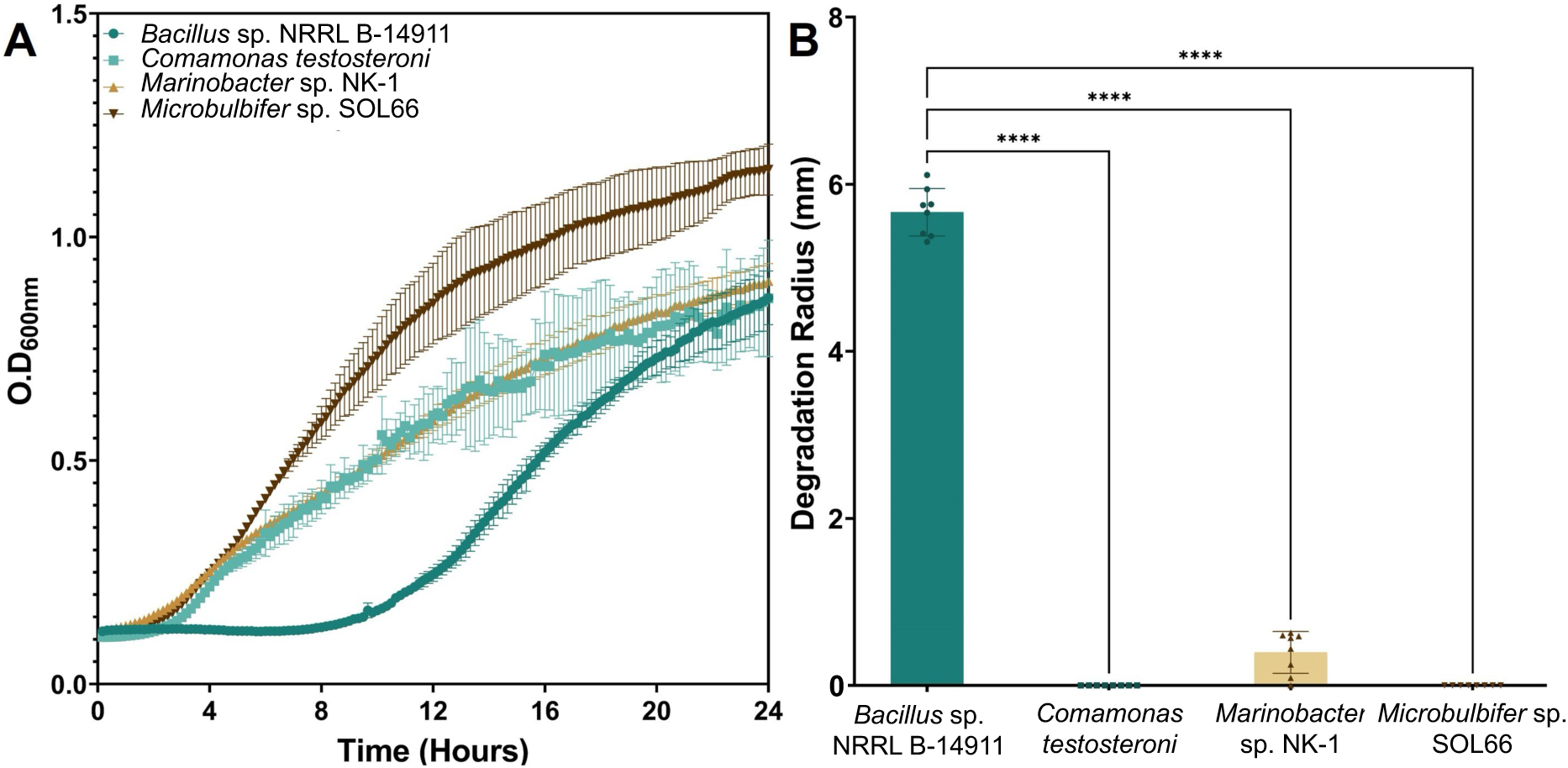
Growth and PHB degradation of marine bacteria. (A) Growth curves of *Bacillus* sp. NRRL B-14911 (dark green), *Comamonas testosteroni* (light green), *Marinobacter* sp. NK-1 (light brown), and *Microbulbifer* sp. SOL66 (dark brown) cultured in Marine Broth at 30 °C. (B) Degradation of PHB by the marine bacteria species was determined by growing bacterial colonies on Marine Broth-agar plates containing PHB powder and measuring the radii of the clear zones of depolymerized PHB formed around the colonies after 7 days (n=8). **** P≤0.0001 by one-way ANOVA statistical analysis To measure the ability of these strains to degrade PHB, cultures of each of the strains were grown on Marine Broth-agar plate containing powdered PHB. A clear zone assay was employed to measure the PHB-depolymerizing activity of these bacterial colonies. Depolymerization of the opaque PHB creates a clear zone within the PHB plate, the radius of which can expand over time at different rates correlating to depolymerization rates (Figure S1).^[19]^ Among the marine bacteria tested, only *Marinobacter* sp. NK-1 and *Bacillus* sp. NRRL B-14911 created measurable clear zones of PHB degradation after 7 days, and the clear zones created by the *Bacillus* sp. NRLL B-14911 colonies were significantly larger than observed for the other strains (Figure 2B). Due to its higher rate of PHB degradation on ocean-mimicking media, *Bacillus* sp. NRLL B-14911 was used throughout the rest of our work.

Next, we evaluated the growth of *Bacillus* sp. NRLL B-14911 in a variety of marine-like culturing conditions. Growth curves of *Bacillus* sp. NRLL B-14911 in Marine Broth at a range of temperatures indicated robust growth of this bacteria at both 30 °C and 25 °C, with somewhat slower growth and lower carrying capacity at 20 °C and 18 °C (Figure S2A). The salt tolerance of *Bacillus* sp. NRLL B-14911 was investigated by measuring growth curves of the strain in liquid growth media with a range of concentrations of added sodium chloride. Robust growth was observed for sodium chloride concentrations ranging from 3-7%, with moderate growth inhibition at 1% sodium chloride (Figure S2B). These results indicate that *Bacillus* sp. NRLL B-14911 shows viability and strong growth in temperatures corresponding to the warmer average surface ocean temperatures, as well as in salinities corresponding to the mean ocean salinity, which is 3.5%.^[20]^

To determine whether 3D-bioprinting parameters including CaCl_2_ crosslinker and alginate bio-ink can affect *Bacillus* sp. NRLL B-14911, we tested its growth and viability under a range of 3D-bioprinting conditions. Ca^2+^ crosslinker concentration can be varied to tune the degree of polymerization of alginate-based hydrogels.^[21]^ Growth curves of *Bacillus* sp. NRLL B-14911 in Marine Broth containing a range of supplemental CaCl_2_ concentrations indicated that bacterial growth could be observed at concentrations between 0.01-0.10 M CaCl_2_ (Figure S2C). Similarly, *Bacillus* sp. NRLL B-14911 bio-printed onto Marine Broth-agar surfaces containing a range of CaCl_2_ concentrations and incubated for 7 days showed no significant change in viability at concentrations from 0.01-0.10 M CaCl_2_ (Figure S2D), well beyond the typical range used for alginate cross-linking in bioprinting applications.^[12a, 22]^ The viability of *Bacillus* sp. NRLL B-14911 in Marine Broth, measured by colony forming units (CFUs), was steady for the first 4 days of incubation but was observed to decrease by ∼4 orders of magnitude after 5 days of incubation (Figure S2E). In contrast, incubation of *Bacillus* sp. NRLL B-14911 in either liquid or polymerized bio-ink resulted in constant high levels of viability over 28 days of incubation (Figure S2E), indicating a protective effect of alginate on bacterial viability. Logistic growth curve modeling of CFU viability data from 3D-bioprinted *Bacillus* sp. NRLL B-14911 predicted a stable plateau phase for 3D-bioprinted bacteria (Figure S2F), suggesting longer-term viability of biomass within bio-stickers that could support extended PHB biodegradation.

### 2.2 Degradation of PHB sheets by 3D-bioprinted bio-stickers

To characterize the PHB-degradation activity of our bio-stickers, *Bacillus* sp. NRLL B-14911 was 3D-printed into bio-stickers that were placed overtop of 1-mm thick PHB sheets and incubated for 28 days at 30 °C (Figure S3). Negative control PHB sheets were incubated with 3D-printed alginate samples that did not contain bacteria. Post-incubation, the bio-stickers were removed, and the cleaned PHB sheets were analyzed by scanning electron microscopy (SEM). The micrographs of the PHB incubated with *Bacillus* sp. NRLL B-14911 bio-stickers showed a corroded morphology characterized by visible pitting, surface irregularities, and microscale roughness (Figure 3A-B). In contrast, the micrographs of the control PHB samples incubated with only 3D-printed alginate showed low surface roughness with little pitting (Figure 3C-D).

**Figure 3.**
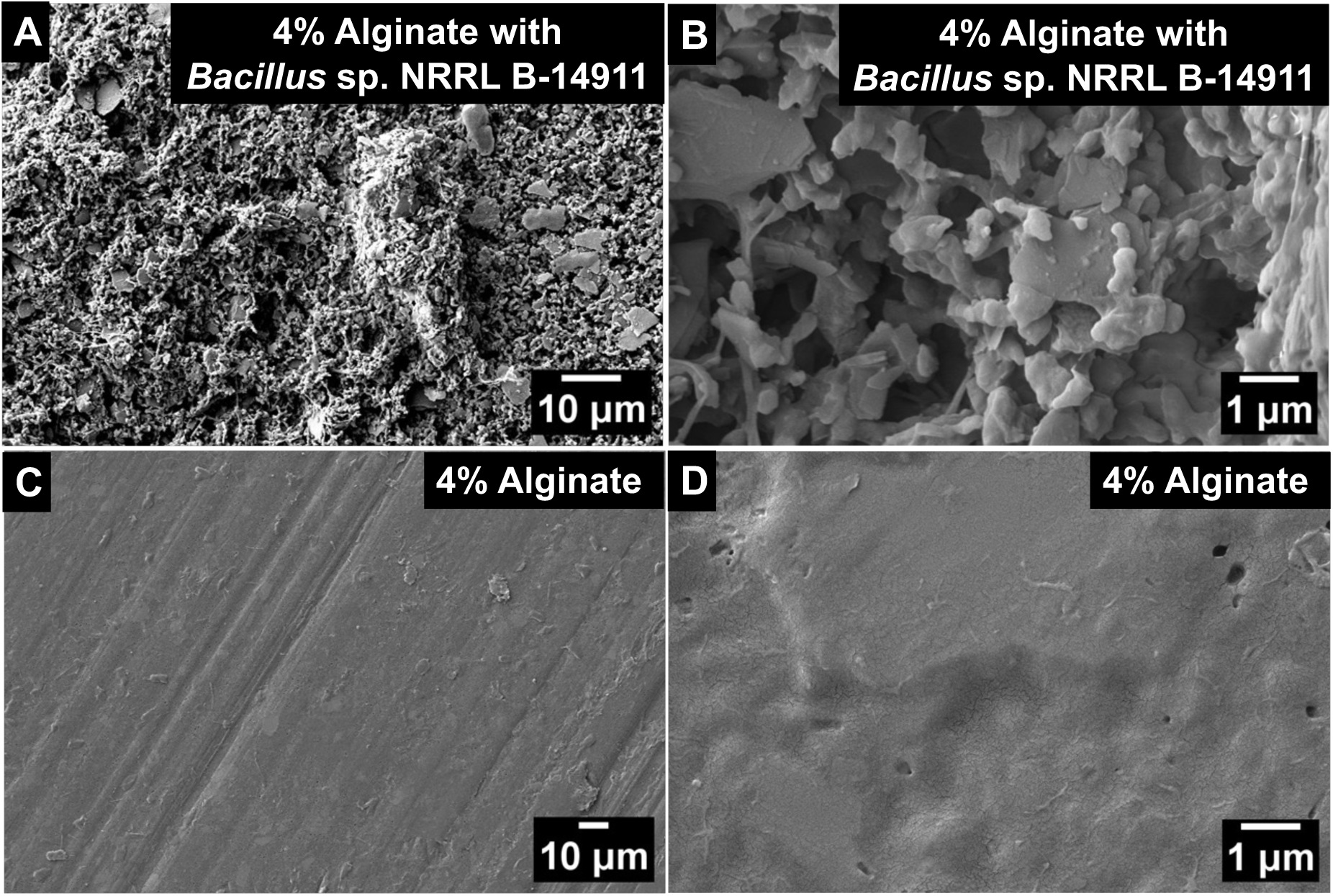
Bio-degradation of solid PHB sheets by 3D-printed *Bacillus* sp. NRLL B-14911. (A,B) SEM images of PHB sheets incubated with a *Bacillus* sp. NRLL B-14911 4% alginate bio-sticker for 28 days. (C,D) SEM images of PHB sheets incubated with 4% alginate bio-stickers not containing bacteria for 28 days. Bio-stickers and bacteria were removed prior to imaging. Micrographs are representative of samples obtained from 3 bio-repeats containing 5 samples each, imaged at multiple locations on the PHB sheets.

To quantify the degradation of solid PHB sheets by 3D-printed *Bacillus* sp. NRLL B-14911 over time, mass loss tests were performed. PHB sheets of 1-mm thickness were cut into discs and sterilized, and each disc was fully covered by a bio-sticker containing *Bacillus* sp. NRLL B-14911. The discs were incubated at 30 °C, and their masses were measured weekly over a four-week period. Significant losses in PHB mass were observed during the incubation period, with a total mass loss of approximately 6% after 28 days of incubation (Figure 4). These results indicate that 3D-printed *Bacillus* sp. NRLL B-14911 is able to bio-degrade solid PHB materials over the course of weeks. Interestingly, the rate of mass loss appeared to decrease after 14 days of incubation. This data suggests that PHB degradation could slow or become limited under our experimental conditions by factors such as limited nutrient availability or buildup of bacteria waste products, which would be expected to be less prevalent under natural ocean conditions with higher water circulation.

**Figure 4.**
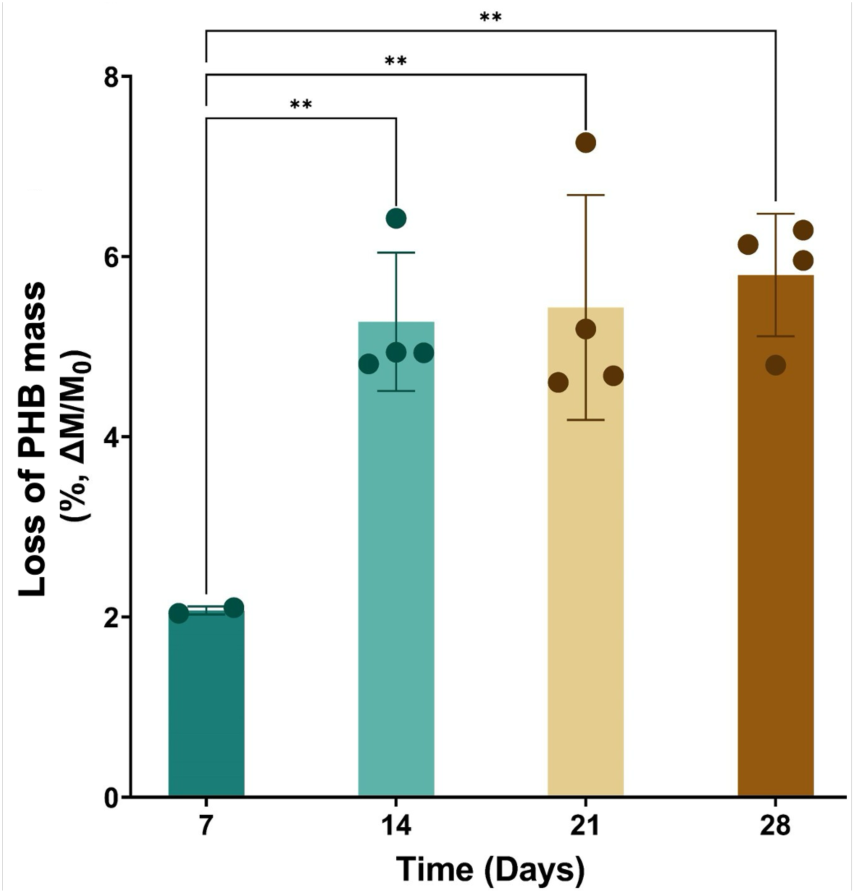
3D-printed *Bacillus* sp. NRLL B-14911 bio-degradation of solid PHB sheets over several weeks. Mass loss of PHB discs incubated with a *Bacillus* sp. NRLL B-14911 4% alginate bio-sticker for 28 days at 30 °C (n=4). ** P≤0.01 by one-way ANOVA statistical analysis.

### 2.3 Tuning of PHB degradation by 3D-bioprinted bio-stickers

To obtain quantitative data on the kinetics of PHB degradation by 3D-printed *Bacillus* sp. NRLL B-14911 bio-stickers, clear zone assays were performed. Bio-stickers were printed onto Marine Broth-agar plates containing PHB powder and calcium chloride to trigger bio-ink gelation and incubated at 30°C. By day 4 of incubation, the regions directly beneath the bio-sticker had been cleared of PHB, and the clear zones continued to expand over 28 days of incubation (Figure 5A). Clear zones within PHB-agar plates could be created by 3D-printed *Bacillus* sp. NRLL B-14911 bio-stickers with a variety of shapes (Figure 5A, Figure S4), indicating that PHB bio-degradation can be carried out by 3D-printed bio-stickers with a range of geometries. Logistic modeling of PHB degradation by 3D-bioprinted *Bacillus* sp. NRLL B-14911 bio-stickers over time indicated that the rate of PHB degradation increased over the first 2-3 weeks of incubation (Figure 5B), predicting sustained and robust PHB degradation ability of bio-stickers over an extended period of time following application. The more sustained PHB degradation observed in these experiments, compared to the mass loss experiments in Figure 4, may be due to the bio-stickers degrading PHB powder, where the smaller particle size can result in more accurate degradation data,^[23]^ as well as the greater ability for secreted PhaZ enzyme to diffuse through the agar plates.

**Figure 5.**
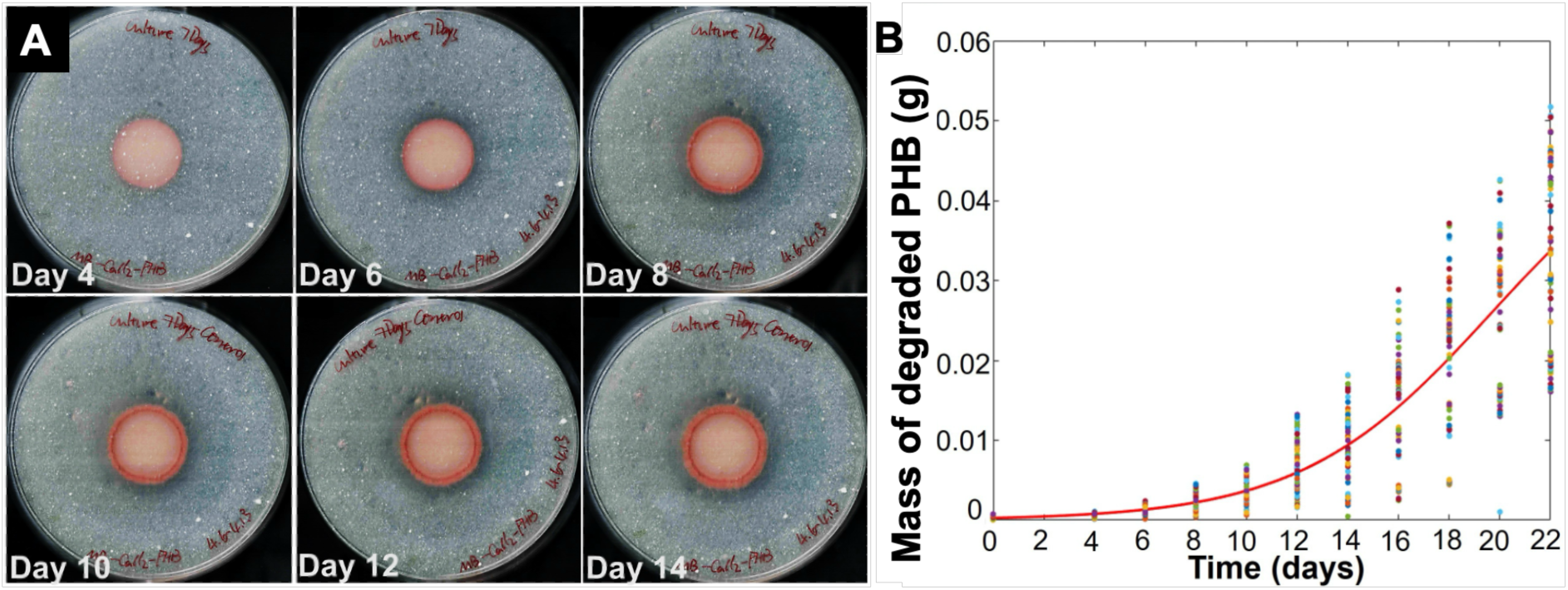
Progression of PHB degradation by 3D-bioprinted bio-stickers. (A) Clear zone assay for *Bacillus* sp. NRRL B-14911 3D-printed into 10-mm diameter circular shapes and placed onto Marine Broth-agar plates containing 0.3 M CaCl_2_ and 0.5% PHB powder. Samples were incubated at 30 °C for 28 days (n=4). (B) The mass of PHB degraded by 3D-printed *Bacillus* sp. NRRL B-14911 over 22 days of incubation was calculated and fit to a logistic equation.

To determine how the PHB bio-degradation rate of our bio-stickers can be tuned, we performed clear zone assays while individually altering several of the experimental parameters. To determine the effect of changing PHB concentration, *Bacillus* sp. NRLL B-14911 bio-stickers were 3D-bioprinted onto Marine Broth-agar plates containing PHB at concentrations ranging from 0.1% – 2.0% and incubated at 30 °C for 22 days. CFU analysis of the bio-stickers at 7-day increments indicated that the concentration of colony-forming units within the bio-stickers decreased by roughly 1-2 logs over the 22-day incubation period but retained a robust 10^7^ CFU/mL after 22 days for the majority of the PHB concentration conditions, including the two conditions with the highest PHB concentrations (Figure S5A). Clear-zone analysis showed significantly higher rates of PHB clear zone expansion at lower PHB concentrations, indicating a negative correlation between PHB concentration and rate of clear zone expansion (Figure 6A-B). The similar CFU values for the bio-stickers over time across all PHB concentrations indicated that the differences in rates of clear zone expansion were not due to changes in the viability of the bio-printed microbes at different PHB concentrations.

**Figure 6.**
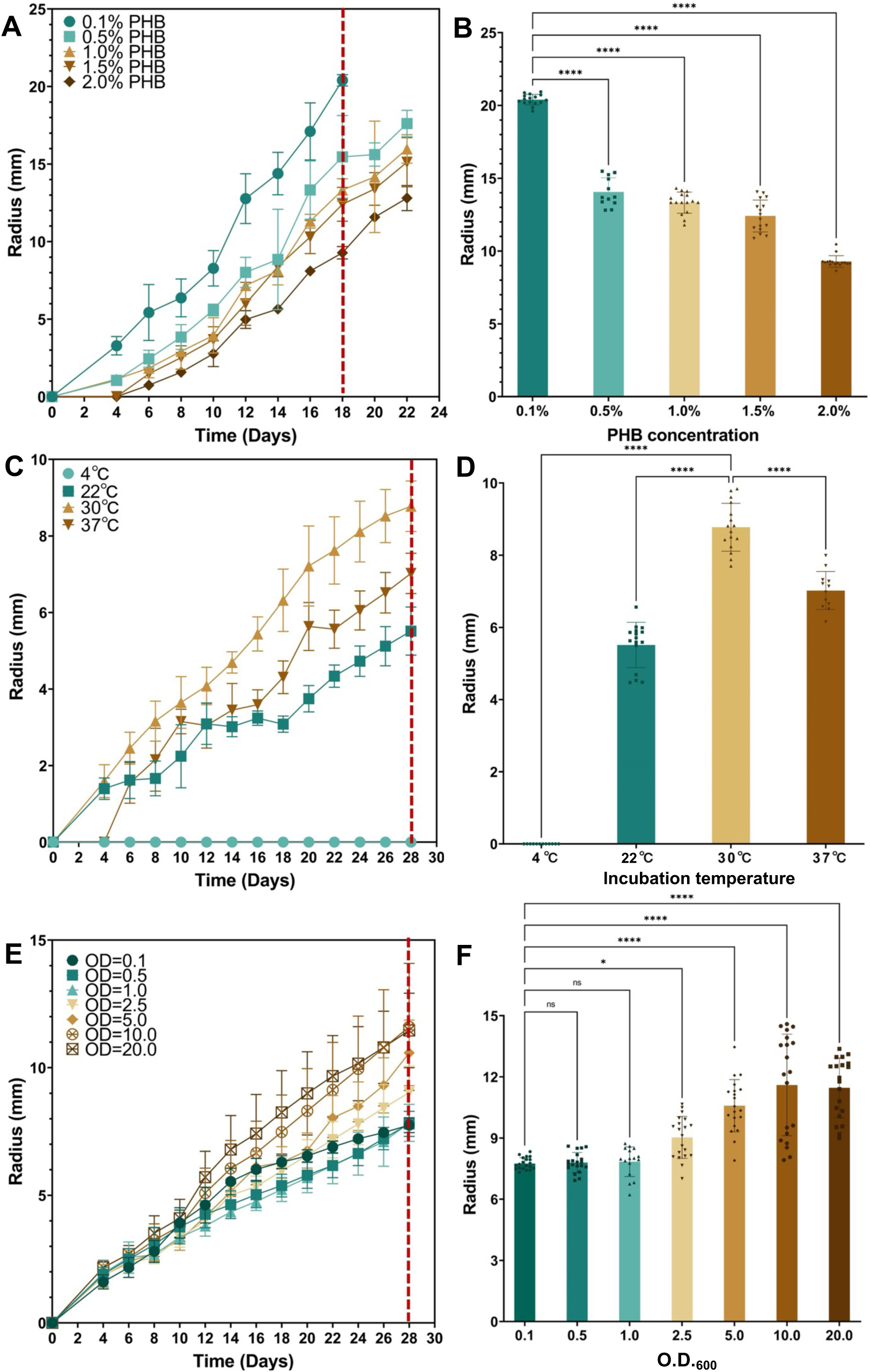
PHB degradation can be tuned by altering bio-sticker parameters. Clear zone assays for *Bacillus* sp. NRRL B-14911 3D-printed at an O.D._600_ of 10 into 10-mm diameter circular shapes and placed onto Marine Broth-agar plates containing 0.3 M CaCl_2_ and 0.5% PHB powder, followed by incubation at 30 °C, except where altered parameters are noted. (A-B) Clear zone radius over time (A) and on day 18 (B) for bio-stickers bio-printed onto plates containing varying PHB concentrations (n=16). (C-D) Clear zone radius over time (C) and on day 28 (D) for bio-stickers incubated at varying temperatures (n=16). (E-F) Clear zone radius over time (E) and on day 28 (F) for bio-stickers printed using bio-ink with varying initial O.D._600_ (n=16). Vertical dashed red lines in panels A, C, and E indicate time points that were analyzed in panels B, D, and F for statistical differences between conditions. * P≤0.05, **** P≤0.0001, ns = not significant by one-way ANOVA statistical analysis

To determine the effect of incubation temperature on bio-sticker PHB degradation efficiency, *Bacillus* sp. NRLL B-14911 bio-stickers were 3D-bioprinted onto Marine Broth-agar plates containing 0.5% PHB and incubated at 4 °C, 22 °C, 30 °C, or 37 °C for 28 days. CFU analysis of the bio-stickers indicated that the concentration of colony-forming units within the bio-stickers stayed roughly constant during the 28-day incubation period at 30 °C, whereas it decreased by roughly 1-2 logs at incubation temperatures of 4 °C, 22 °C, or 37 °C (Figure S5B). Clear-zone analysis showed that PHB degradation by the bio-stickers was fastest at 30°C, significantly slower at 22 °C and 37 °C, and not detectable at 4 °C (Figure 6C-D). The rate of PHB degradation at 37 °C increased after the first 6 days of incubation, after which it was similar to the degradation rate at 30 °C, perhaps indicating adaptation to the higher temperature condition over time. The lack of detectable PHB degradation at 4 °C despite the high viability of the cells within the bio-sticker indicates that the bacteria at this colder temperature remain alive but with very low or nonexistent PHB degradation activity, showing that microbial viability and PHB degradation activity are not necessarily linked.

To determine the effect of the concentration of initial biomass on bio-sticker PHB degradation efficiency, *Bacillus* sp. NRLL B-14911 bio-stickers were 3D-bioprinted using bio-ink with cell densities varying from O.D._600_ 0.1 to O.D._600_ 20 onto Marine Broth-agar plates containing 0.5% PHB and incubated at 30 °C for 28 days. CFU analysis of the bio-stickers showed that the concentration of colony-forming units within the bio-stickers was approximately10^8^ CFU/mL at the 7-day timepoint for all samples and exhibited a consistent gradual decrease by 1-2 logs over the incubation period, regardless of initial cell density (Figure S5C). The similar CFU/mL values of the bio-stickers at the 7-day timepoint may indicate that the 3D-bioprinted bacteria were able to replicate up to a comparable carrying capacity for these specific incubation conditions during the first week after deposition. Clear-zone analysis indicated that higher initial biomass corresponded to significantly greater PHB degradation over time, with the highest-O.D. bio-inks (e.g. O.D._600_ 10, 20) exhibiting clear zones with approximately 65% larger radius after 28 days compared to the lowest-O.D. bio-ink (O.D._600_ 0.1) (Figure 6E-F). These results indicate that initial cell density within bio-stickers enhances PHB degradation efficiency, despite the cell viability within the bio-stickers remaining roughly similar throughout days 7-28 of the experiment, potentially due to increased accumulation of PHB-degrading enzymes during early time periods within the experiment.

While changes in PHB concentration, incubation temperature, and initial biomass were all observed to robustly tune the rate of PHB degradation by 3D-bioprinted bio-stickers, altering other 3D-bioprinting parameters had less effect on PHB degradation rate. To test the effect of bio-sticker hydrogel density on PHB degradation, *Bacillus* sp. NRLL B-14911 bio-stickers were 3D-bioprinted using bio-ink with alginate concentrations varying from 2% - 6% onto either Marine Broth-agar plates or PBS-agar plates containing 0.5% PHB and incubated at 30 °C. PHB degradation rates varied by less than 15% (Figure S6A-B) and CFU/mL values differed by less than 1 log (Figure S6C) across all alginate concentrations. To test the effect of bio-sticker height on PHB degradation, *Bacillus* sp. NRLL B-14911 was 3D-bioprinted into bio-stickers with heights ranging from 1-15 mm onto Marine Broth-agar plates containing 0.5% PHB and incubated at 30°C. Most bio-stickers of varying heights showed statistically indistinguishable differences in PHB degradation, with a modest increase for the tallest bio-stickers (Figure S6D-E). CFU/mL values were uniformly high across all bio-sticker heights, with the exception of the shortest 1-mm tall bio-stickers (Figure S6F), which may have lost viability more rapidly due to higher susceptibility to dehydration. To test the effect of bio-sticker radius on PHB degradation, *Bacillus* sp. NRLL B-14911 was 3D-bioprinted into bio-stickers with radii ranging from 10-25 mm onto Marine Broth-agar plates containing 0.5% PHB and incubated at 30 °C. PHB degradation was similar across bio-stickers of all radii, with a significant but minor 2-mm increase in PHB degradation radius for the smallest 10-mm bio-sticker (Figure S6G-H) and similar high CFU/mL for the bio-stickers that varied by approximately 1 log across a 24-day time period (Figure S6I). These results indicate that PHB degradation rate of the bio-stickers is only modestly affected by their geometry and hydrogel cross-linking density.

### 2.4 3D-bioprinted bio-stickers retain PHB-degradation activity following storage and transfer

For deployment in application scenarios to degrade bioplastic objects, 3D-bioprinted bio-stickers will likely be fabricated and then temporarily stored or transported to end users for utilization. The bio-stickers would need to retain viability and PHB-degradation activity for several days or weeks in order to still be able to depolymerize PHB once transferred to their target objects following a period of storage or transportation. To test this scenario, *Bacillus* sp. NRLL B-14911 bio-stickers were 3D-bioprinted onto Marine Broth-agar plates containing 0.5% PHB and stored for 7 or 14 days, then transferred to a fresh Marine Broth-PHB-agar plate using sterile tweezers to test for PHB-degradation activity (Figure 7A). The bio-stickers were able to be transferred to the new plates without breaking, adhered well to the tweezers throughout the transfer, and adhered to the new plate post-transfer (Video S1). Clear-zone analysis indicated that bio-stickers that had been transferred after 7 or 14 days of storage showed robust PHB degradation (Figure 7B-C), accompanied by uniformly high CFU/mL for the bio-stickers across all conditions (Figure 7D). Transferred bio-stickers showed higher PHB degradation at the earliest time points, likely due to the printed microbes having had additional time during the storage period to divide and accumulate biomass (Figure 7B). After a delayed burst of PHB degradation, the non-transferred bio-stickers adopted a final rate of PHB degradation that was similar to the transferred bio-stickers (Figure 7B), indicating that the bio-stickers can maintain a steady-state rate of PHB degradation for several weeks post-printing.

**Figure 7.**
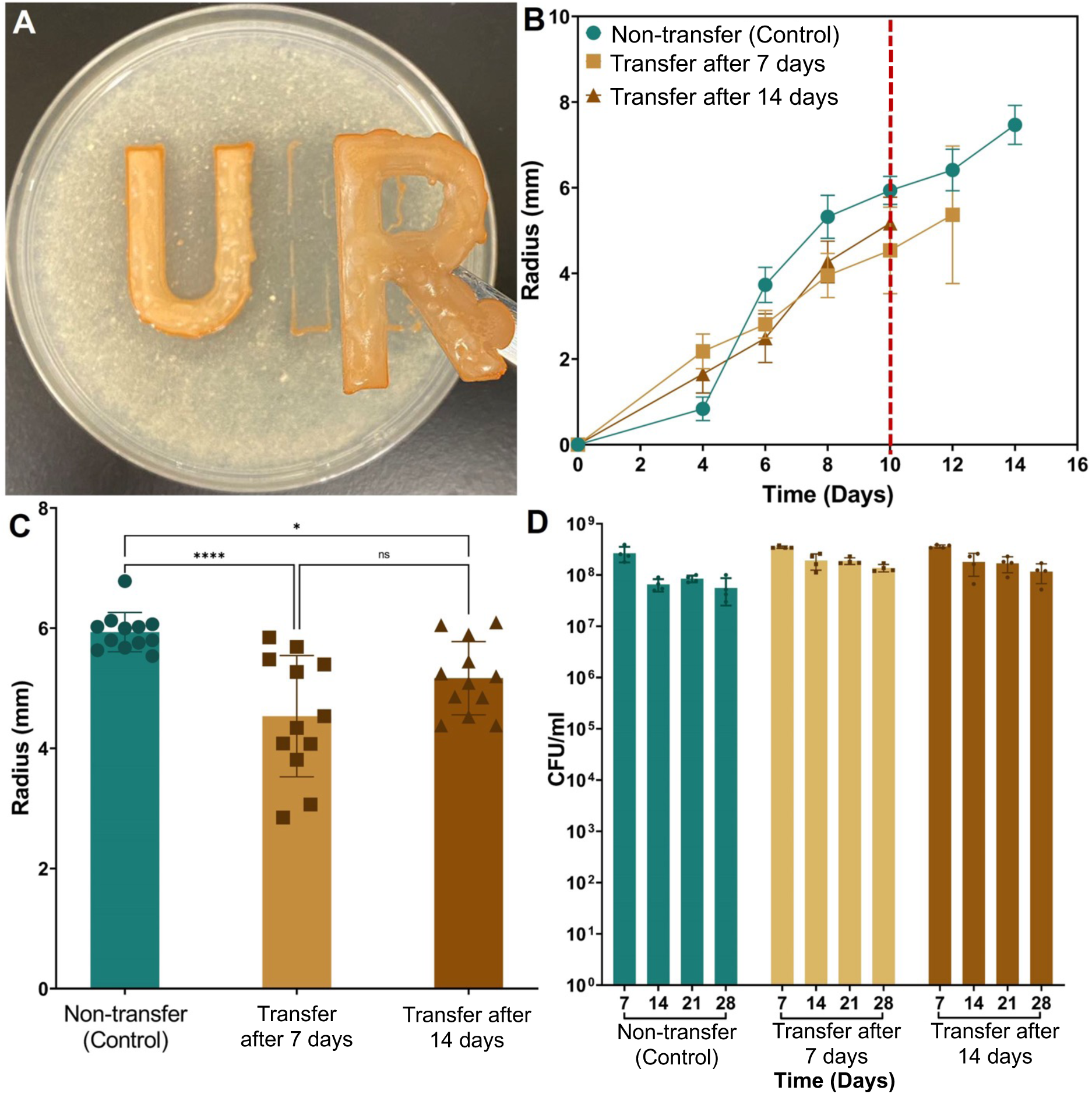
Transferred bio-stickers retain PHB-degradation activity. (A) *Bacillus* sp. NRRL B-14911 bio-stickers can be transferred following incubation on Marine Broth-PHB-agar plates. (B-C) Clear zone radius over time (B) and on day 10 (C), and CFU/mL over time (D) for bio-stickers bio-printed onto Marine Broth-PHB-agar plates (green) or transferred onto fresh Marine Broth-PHB-agar plates following 7 (tan) or 14 (brown) days of storage at 30°C (n=16). Vertical dashed red line in panels B indicates time points that were plotted in panel C. * P≤0.05, **** P≤0.0001, ns = not significant by one-way ANOVA statistical analysis

### 2.5. Mechanical properties of 3D-bioprinted bio-stickers

Since PHB-degrading bio-stickers will need to maintain their mechanical robustness over several weeks of storage and application to target PHB objects, we tested their mechanical properties in a range of environmental conditions. To test the viscoelastic behavior of the 3D-bioprinted hydrogels, rheology was performed. Bio-stickers either containing *Bacillus* sp. NRLL B-14911 or not containing microbes in 4% alginate were stored on Marine Broth-agar plates containing 0.5% PHB for 21 days at either 4 °C or 30 °C, to simulate temperatures towards the higher and lower range of typical ocean water temperatures. Strain sweep oscillatory tests were performed on the bio-stickers to determine their storage modulus (G’) and loss modulus (G’’) (Figure 8A, Figure S7). Notably, for all sample groups, at strains less than 1%, G’ was consistently higher than G’’ by approximately one order of magnitude, indicating that the samples exhibited predominantly elastic, gel-like behavior. The storage modulus G’ was approximately 10^5^ Pa for the samples stored at 4 °C either containing or not containing microbes, as well as the samples without microbes stored at 30 °C, which is a typical value for alginate-based materials.^[24]^ In contrast, the samples containing *Bacillus* sp. NRLL B-14911 stored at 30 °C had a lower storage modulus around 10^4^ Pa (Figure 8A). The samples containing *Bacillus* sp. NRLL B-14911 and stored at 30 °C reached the crossover point, where G’ intersects with G’’, at a significantly lower strain compared to the other groups (Figure 8B). These samples also displayed lower G’ and G’’ modulus levels in the linear viscoelastic (LVE) region compared to the other groups (Figure 8C). These results indicate a more compliant gel structure in the presence of microbes after storage at 30 °C, potentially due to the bacteria cells weakening the crosslinking density and structural stability through its biodegradation activity. Interestingly, the samples stored at 4 °C did not display any loss of stiffness or viscoelasticity with the presence of microbes, which could indicate that storing the bio-stickers at cooler temperatures reduces bacteria-driven biodegradation of the alginate hydrogel matrix.

**Figure 8.**
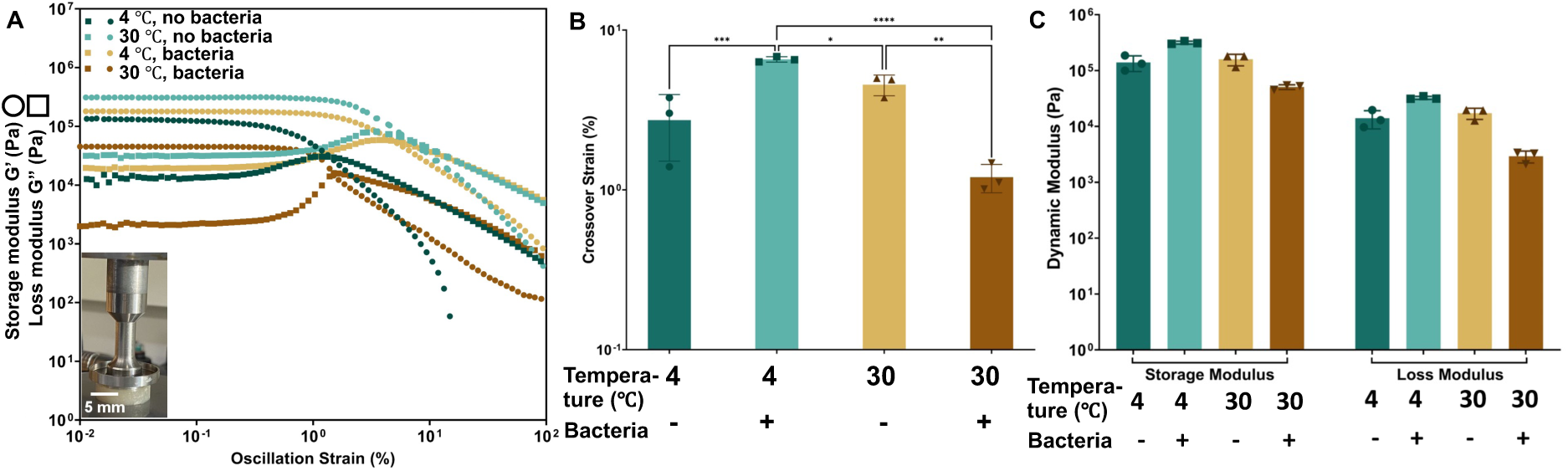
Rheological analysis of bio-stickers indicates viscoelastic behavior. (A) Dynamic moduli plotted against oscillation strain for 3D-printed bio-stickers including or not including *Bacillus* sp. NRRL B-14911 and incubated at either 4°C or 30°C for 21 days. The inset image depicts the equipment setup used for testing. (B) Crossover strain values for each type of sample, indicating the critical strain where storage and loss moduli intersect (n=3). (**C**) Storage and loss modulus values for samples in the LVE region (n=3). * P≤0.05, ** P≤0.01, *** P≤0.001, **** P≤0.0001 by one-way ANOVA statistical analysis

To assess the tensile strength, stiffness, and toughness of the bio-stickers, we performed monotonic tensile tests to demonstrate the material’s response to uniaxial tension. Bio-stickers either containing *Bacillus* sp. NRLL B-14911 or not containing microbes in 4% alginate were 3D-bioprinted into “dog-bone” structures following the ASTM D638-05 standard.^[25]^ The bio-stickers were incubated on Marine Broth-agar plates containing 0.5% PHB for 21 days at either 4 °C or 30 °C prior to tensile testing (Figure 9A). The stress-strain curves for each of the bio-sticker samples displayed an initial region in which the stress increased proportionally with strain, followed by a decrease in slope, and then fracture (Figure 9B; Figure S8). The modulus of elasticity, which characterizes the material’s stiffness, was determined from the slope of the stress-strain curve for each material. The elastic modulus for each of the bio-sticker samples was around 1MPa (Figure 9C). No significant differences in elastic modulus were seen when the bio-stickers were stored at 4 °C versus 30 °C, or for bio-stickers with or without *Bacillus* sp. NRLL B-14911. The ultimate tensile strength (UTS) of the samples was determined as the maximum tensile load that the materials could endure before failure. The bio-sticker samples incubated at 30 °C, both with and without microbes, demonstrated significantly higher UTS values than the bio-stickers incubated at 4 °C (Figure 9D). The inclusion of bacteria in the bio-stickers resulted in a significant decrease in UTS for bio-stickers incubated at 30 °C but not at 4 °C (Figure 9D), potentially due to enzymatic bio-degradation of the alginate matrix by the microbes at the higher temperatures.

**Figure 9.**
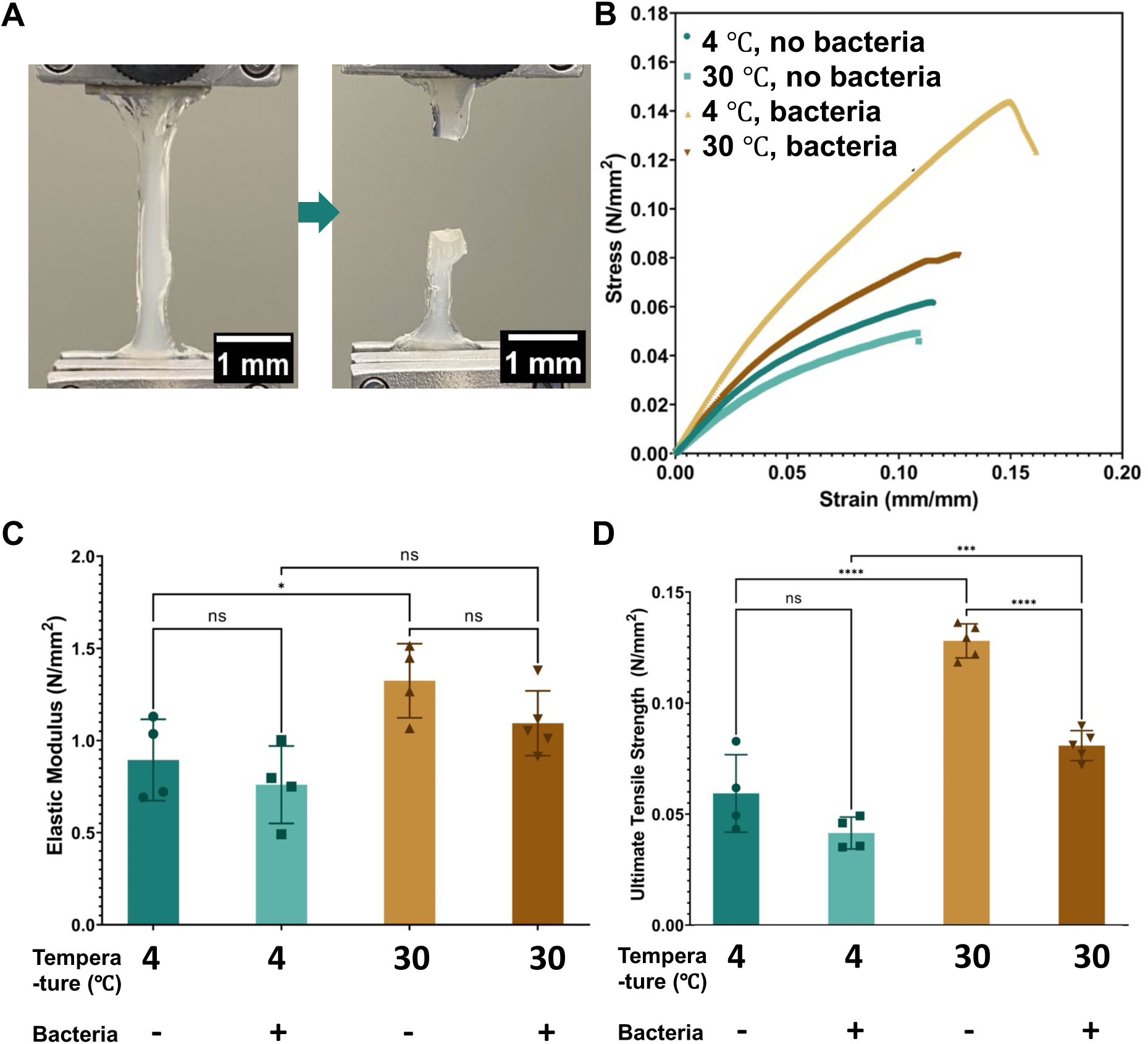
Tensile testing of bio-stickers. (A) Image of a 3D-printed bio-sticker undergoing a tension loading test. (B) Strain-stress curves for 3D-printed bio-stickers including or not including *Bacillus* sp. NRRL B-14911 and incubated at either 4 °C or 30 °C for 21 days. (C) The elastic modulus for each type of sample and (D) the ultimate tensile strength for each type of sample (n=4-5). * P≤0.05, *** P≤0.001, **** P≤0.0001, ns = not significant by one-way ANOVA statistical analysis

In order to efficiently biodegrade PHB objects, bio-stickers will need to adhere to PHB materials to keep their embedded PHB-degrading microbes in close proximity to the target bioplastics. To assess the adhesion strength of bio-stickers when bonded to PHB plates, adhesion tests were conducted. Bio-stickers either containing *Bacillus* sp. NRLL B-14911 or not containing microbes in 4% alginate were 3D-bioprinted into rectangular structures and incubated on Marine Broth-agar plates for 21 days at either 4 °C or 30 °C prior to adhesion testing (Figure 10A). Inspired by lap shear testing protocols,^[26]^ PHB plates and bio-sticker samples were positioned in opposite grips of a universal testing machine with an overlap distance of 20 mm. The PHB and bio-stickers were placed into contact through gentle pressure, then the applied force was measured while the PHB and bio-stickers were vertically displaced from each other at a rate of 1 mm/s (Figure 10B; Figure S9). The maximum load value, representing the peak force that the bio-stickers could endure before separating from the PHB surface, is directly proportional to the adhesion strength. Our data indicated that the average maximum load values for all samples fell within the range of 0.15-0.2 N (Figure 10C). The energy dissipation value, calculated from the area under the force-displacement curve, indicates the energy expended by the bio-sticker until interface failure. An increased energy dissipation capacity is typically associated with more durable and dependable adhesion under diverse conditions.^[27]^ Our results showed that average energy dissipation values for all sample types ranged between 0.09-0.17 mJ (Figure 10D). Notably, the presence or absence of the *Bacillus* sp. NRLL B-14911 and the variations in incubation temperature did not have any significant effects on the measured adhesive properties of the bio-stickers (Figure 10C-D).

**Figure 10.**
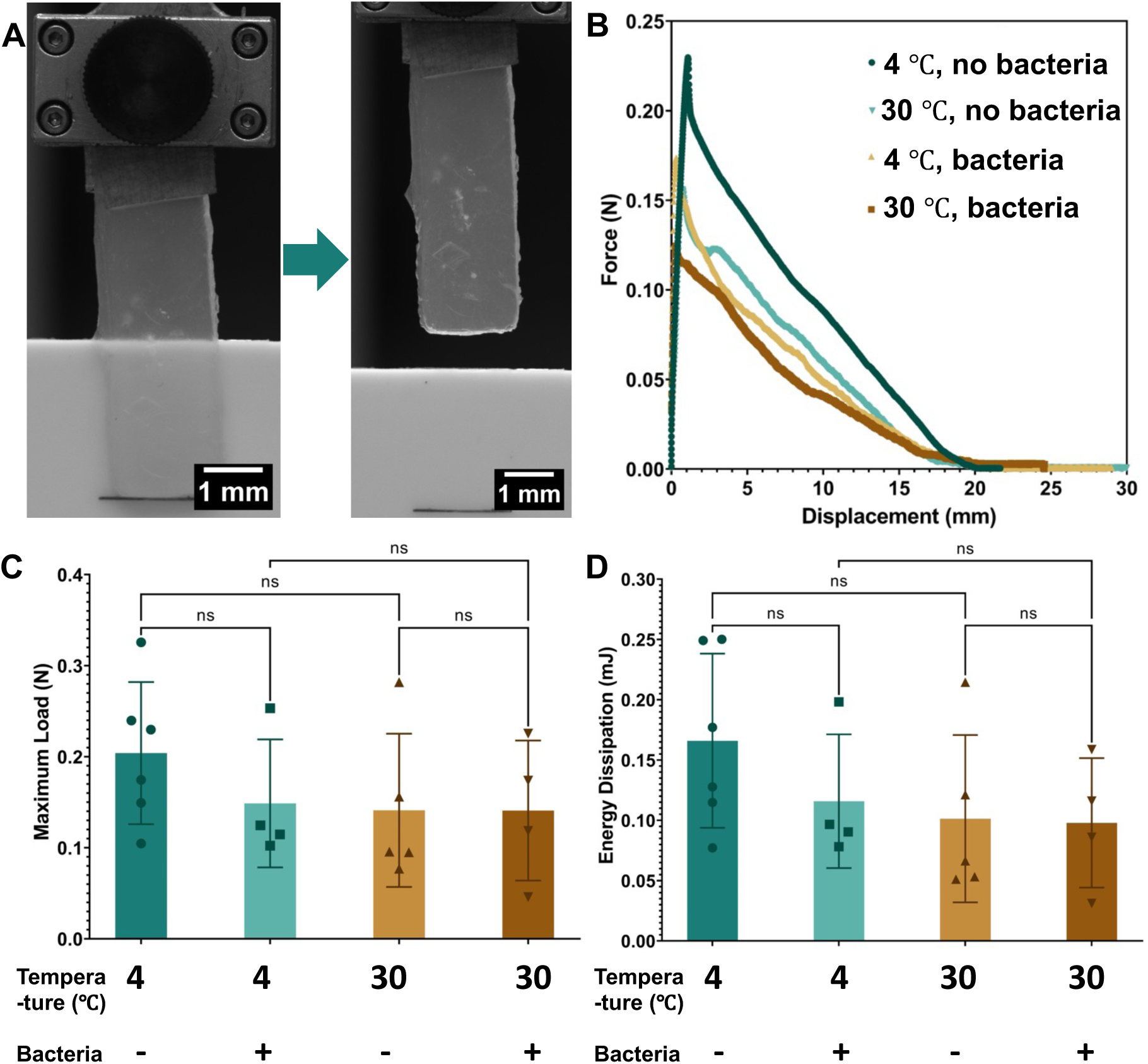
Adhesion testing of bio-stickers. (A) Image of a 3D-printed bio-sticker undergoing an adhesion test. (B) Force-displacement curves for 3D-printed bio-stickers including or not including *Bacillus* sp. NRRL B-14911 and incubated at either 4 °C or 30 °C for 21 days. (C) The maximum load for each type of sample and (D) the energy dissipation for each type of sample (n=4-6). ns = not significant by one-way ANOVA statistical analysis

## 3. Conclusion

In this study, we have developed a 3D-bioprinted “bio-sticker,” consisting of marine bacteria integrated within an alginate hydrogel, for accelerating and tuning the degradation of PHB bioplastics in marine environments. The alginate hydrogel provides a supportive scaffold for the bacteria that spatially distributes the bacteria^[22d]^ and localizes them to the surface of a target PHB object, ensuring that secreted PHB-depolymerizing enzymes are delivered to the substrate. The method of 3D-bioprinting of the bio-stickers offers customizable shapes and dimensions,^[28]^ enabling a user to pattern the living material onto target objects of varied geometries. This versatility is especially relevant for marine devices that might require selective or patterned degradation over specific areas, as opposed to uniform, bulk material degradation.

Our results demonstrate the tunability of PHB biodegradation through altering and optimizing bio-sticker parameters. By adjusting initial bio-sticker cell density, PHB concentration, and incubation temperature, the rate of PHB degradation can be significantly changed. Higher initial bacterial biomass in the bio-stickers yielded faster PHB depolymerization, a parameter can that be altered in a straightforward manner during bio-ink preparation. Additionally, cooler temperatures were seen to significantly slow the rate of PHB degradation, likely through decreasing microbial metabolism. Thus, the specific local climate and current weather conditions experienced on-site following deployment of a PHB-degrading bio-sticker may either shorten or extend the lifetime of the target plastic object prior to eventual breakdown. In practical terms, these findings offer a route to customize biodegradation rates for diverse marine applications, ranging from short-lived devices that require rapid disposal, to robust sensors and buoys that must function over extended periods prior to end-of-life.

Mechanical testing of the bio-stickers revealed that the mechanical integrity of the bio-stickers at lower temperatures remains largely unaffected by the incorporation of microbes. Specifically, at 4 °C the presence of bacteria within the bio-stickers did not compromise the hydrogel’s rheological properties, elastic modulus, ultimate tensile strength, or adhesion to PHB, suggesting that the bio-stickers can likely be stored or deployed under cooler oceanic conditions without significant mechanical breakdown of either the hydrogel or the substrate. In contrast, while the PHB-degrading microbe employed in this study exhibited the highest rate of PHB degradation at 30 °C, incubation of bio-stickers at this temperature was associated with a more pronounced impact on the hydrogel’s mechanical integrity. The presence of microbes within bio-stickers incubated at 30 °C did not change the elastic modulus or adhesion to PHB but did result in lower storage and loss moduli as well as ultimate tensile strength, likely due to higher metabolic activity of the embedded microbes. Given that many regions of the ocean are cooler than 30 °C, future work will entail identifying or engineering bacterial strains capable of high rates of PHB biodegradation at lower temperatures, possibly by leveraging psychrotolerant or deep-sea microbes. Nevertheless, our findings show that the salt tolerance of *Bacillus* sp. NRRL B-14911 is sufficient for typical ocean salinities, suggesting that salinity may be less of an obstacle than temperature in adapting these bio-stickers to specific use-cases.

From an application standpoint, 3D-bioprinted PHB-degrading bio-stickers could be straightforwardly applied to a variety of marine-deployable instruments and objects to accelerate their biodegradation in ocean environments. Bio-stickers could be used on disposable or single-use sensing devices or objects. Another potential application lies in the emerging sphere of the Internet of Things (IoT) and autonomous ocean sensing, where devices are increasingly used for extended monitoring and data collection but are not retrieved after deployment into the ocean.

For example, CARTHE surface drifters are deployed by the hundreds to track ocean currents but are typically only monitored for a few weeks after release.^[29]^ Additionally, the National Oceanic and Atmospheric Administration’s Global Drifter Program and Argo float program each employ hundreds or thousands of devices and are examples of expendable ocean platforms that could be made biodegradable. Incorporating biodegradable PHB plastic structural components plus a user-applied bio-sticker just prior to ocean deployment would allow such devices to biodegrade after their operational lifetime, thereby decreasing their lifetimes and reducing plastic waste in their target marine environments. The 3D-printed bio-sticker format ensures that the bio-stickers can be conformed to the shape of the device, delivering on-demand or location-specific degradation.

Newly developed methods to accelerate plastic degradation rates are increasingly using biologically-driven approaches. Recent works have created plastic materials that contain bacterial spores that act as living fillers and can survive the high-temperature process of plastic forming to facilitate degradation in compost or soil environments at end-of-life.^[30]^ Additionally, there are several notable examples of integrating polymer-degrading enzymes into plastics, often together with stabilizing molecules that preserve enzymatic activity both during plastic molding or extrusion and within the resultant solid-state plastic environment.^[31]^ In contrast, our approach of applying the PHB-degrading microbes within a bio-sticker onto a fully formed plastic object ensures that the microbes and their PHB-depolymerizing enzymes can be kept in aqueous environments that support their activity and viability. Furthermore, the room-temperature 3D-bioprinting process prevents the need for the plastic-degrading microbes and their enzymes to undergo elevated temperatures and/or pressures associated with the various techniques for molding or 3D-printing plastic objects. Since the bio-sticker’s microbes do not need to be incorporated within the plastic objects at the time of their production, any adverse effects on the mechanical properties of the plastic materials that arise from embedded microbes or enzymes can be avoided, and there are no effects on the product’s shelf life that may result from extended exposure to the embedded microbes or enzymes. Also, an applied bio-sticker can be removed or replaced partway through the biodegradation process, providing more control over the dynamics of the plastic breakdown over time and allowing for flexibility in redesigning or reusing the plastic substrate.

Overall, this work demonstrates a scalable and tunable strategy to mitigate ocean debris while integrating eco-friendly materials into the rapidly expanding blue economy. By addressing the inherent challenges to plastic degradation posed by marine environments, our 3D-bioprinted bio-stickers can help transform the fate of bioplastic objects, enabling responsible disposal without sacrificing their original performance. The potential to extend these methods to other biodegradable plastic polymers, as well as to apply or engineer new microbial strains with enhanced bioplastic degradation activity in a range of ocean-relevant conditions offers a wealth of directions for future research. Ultimately, these advancements can bolster autonomous marine sensing and IoT deployments in ways that greatly reduce long-term ecological impact.

## 4. Experimental Methods

### Bacteria strains and culturing conditions

The bacteria strains used in this study are *Bacillus* sp. NRRL B-14911^[14–15]^ (Agricultural Research Service Culture Collection, United States Department of Agriculture), *Comamonas testosteroni*^[16]^ (ATCC 11996), *Marinobacter* sp. NK-1^[17]^ (ATCC 700491), and *Microbulbifer* sp. SOL66^[18]^ (ATCC 70072). Cultures of bacteria were grown in Marine Broth 2216 medium (Merck KGaA, Darmstadt, Germany) under continuous shaking at 220 rpm at 30 °C to an optical density (O.D._600_) of 1.0. A 50 mL aliquot of the cultures were centrifuged at 1500 rpm for 20 minutes, and the resulting bacterial pellet was resuspended in either bio-ink (4% alginate in Marine Broth) or Marine Broth. Resuspended cultures were incubated at 30 °C with shaking for 48 hours. Samples removed at 0 and 48 hours were subjected to CFU assays.

### >Growth curves

Cells were cultured as described above, with the exception that cultures were grown in a 96-well plate. The plate was incubated at 30 °C with continuous orbital shaking at 220 rpm in a microplate reader (BioTek Synergy H1), which recorded O.D._600_ measurements every 15 minutes for a total of 24 or 48 hours.

### Colony forming unit assays

For liquid cultures, cells were resuspended to an O.D._600_ of 0.1 in Marine Broth and were serially diluted in 1X PBS. For bio-stickers, the bio-sticker was dissolved in an equal volume of 0.1 M CaCl_2_, and resuspended cells were serially diluted in 1X PBS. From each dilution, a 100 µL aliquot was plated onto Marine Broth-agar plates. Plates were incubated overnight at 30° C, and colonies were counted the following morning. The following equation was used to calculate the CFU/mL of the samples: (number of colonies*dilution factor)/volume of cells plated.

### 3D Bioprinting

Bio-inks for 3D-bioprinting were prepared by growing an overnight culture of *Bacillus* sp. NRRL B-14911 in Marine Broth at 30 °C, harvesting the cells by centrifugation at 1500 rpm for 20 minutes, and re-suspending the cells to an O.D._600_ of 10 with a sterile solution of sodium alginate [4% (w/v)] in Marine Broth. Bio-ink was loaded into a sterile syringe and attached to a custom-built 3D-bioprinter.^[12a, 22a]^ Digital models were created using computer-aided design CAD software (Autodesk Fusion 360), which was then sliced using slicing software (Simplify3D). 3D-bioprinting was carried out using print speed of 1-5 mm/s, with bio-ink extrusion rate adjusted to achieve a layer height of 0.25 mm. Unless otherwise specified, bio-stickers were printed using bio-ink with O.D._600_ of 10 in 4% alginate-Marine Broth, to create a bio-sticker with radius of 10 mm and height of 0.25 mm, printed onto Marine Broth-PHB-agar plates (50% Marine Broth, 0.5% PHB (Mango Materials, USA), and 0.3 M CaCl_2_).

### Clear zone assays of PHB degradation

*Bacillus* sp. NRRL B-14911 was harvested, resuspended in 4% alginate-Marine Broth, and 3D-bioprinted onto Marine Broth-PHB-CaCl_2_ agar plates. After solidification, the bio-stickers were transferred to fresh plates if needed using a sterile spatula. Plates were incubated at 30 °C, and PHB degradation was monitored on a daily basis by measuring the size of the clear zone as the average perpendicular distance of the edge of the clear zone away from the edge of the bio-sticker.

### SEM imaging

*Bacillus* sp. NRRL B-14911 was cultured overnight to stationary phase in Marine Broth and bio-printed into bio-stickers using a bio-ink with O.D._600_ of 10 in 4% alginate-Marine Broth, to create bio-stickers with radius of 10 mm and height of 0.25 mm. These bio-stickers were applied to degrade PHB discs (1-mm thickness, 44.5-mm diameter, Mango Materials, USA) over 28 days at 30 °C. Post-incubation, the bio-stickers were removed from the PHB films, which were cleansed and freeze-dried in preparation for SEM analysis. The films were sputter-coated with approximately 5 mm gold film at 20 mA for 60 seconds and positioned onto SEM stubs. SEM imaging was performed on a JEOL JSM-IT500HR InTouchScope^TM^ scanning electron microscope at an accelerating voltage of 5 kV, and backscattered electron images were captured at multiple magnifications.

### Mass loss analysis of PHB discs

PHB discs (1-mm thickness, 44.5-mm diameter) were sterilized with ethanol, and each disc was fully covered by a 3D-bioprinted bio-sticker containing *Bacillus* sp. NRLL B-14911. The discs were incubated at 30 °C. Post-incubation discs were removed from the bio-sticker, cleaned, and weighed to determine mass loss.

### Rheology testing

Bio-stickers were prepared by mixing 4% alginate bio-ink solutions with or without *Bacillus* sp. NRRL B-14911. The bio-ink solutions were cast onto CaCl_2_-Marine Broth agar plates to polymerize the bio-ink into bio-stickers, after which they were incubated at 4 °C or 30 °C for 21 days. Shear rheology was performed using a TA HR-30 rheometer, using a 20-mm parallel plate geometry on a steel Peltier plate to ensure a uniform shear field during testing. Strain sweep tests were performed at a frequency of 1 Hz on a range from 0.01% to 100% strain to determine the limit of the linear viscoelastic range and to identify the material’s yield point. The dynamic moduli were then analyzed, specifically the storage modulus (G’) and loss modulus (G’’).

### Tensile testing

Bio-stickers were 3D-bioprinted using 4% alginate bio-ink with or without *Bacillus* sp. NRRL B-14911 into the ASTM D638-05 standard dogbone shape,^[25]^ with an average sample thickness of 1.528 mm (ranging from 0.724 mm to 2.047 mm). Samples were 3D-bioprinted onto CaCl_2_-Marine Broth agar plates and incubated at 4 °C or 30 °C for 21 days. The samples were washed with 0.1 M CaCl_2_ solution prior to removal. Quasi-static tensile testing was performed via monotonic tensile loading to failure on a universal testing machine (Zwick-Roell Z010) with a 20 N load cell (X-Force HP, Zwick-Roell), at a displacement speed of 1.1 mm*s^-1^. Non-woven fabrics were applied to reduce specimen slippage in the grips. Specimen width and thickness were measured optically, and the gauge length was measured with calipers. Engineering strain was calculated as crosshead displacement divided by the initial gauge length. Engineering stress was calculated as the force divided by the cross-sectional area. Elastic modulus was calculated by fitting a linear regression to the range from 0 to 0.15 strain (mm/mm) of the stress-strain curve and selecting the maximum value. Ultimate stress was reported based on the maximum stress prior to failure for samples that broke in the center section.

### Adhesion testing

Adhesive properties of the bio-stickers were tested via a protocol inspired via lap-shear testing protocols.^[26]^ Rectangular bio-stickers (L*W*H=47.22 mm*12.65 mm*3.71 mm) were 3D-bioprinted using 4% alginate bio-ink with or without *Bacillus* sp. NRRL B-14911 onto CaCl_2_-Marine Broth agar plates and incubated at 4 °C or 30 °C for 21 days. PHB sheets (thickness 1.99 mm, Goodfellow UK) were gripped in a vertical fixed position within a universal testing machine (Zwick-Roell Z010) with a 20 N load cell. The bio-stickers were tightened in the top grip, and the surface patted dry. The bio-stickers were pressed into contact with the PHB sheet with an overlap distance of 18.87 mm. The bio-sticker specimens were pulled at a constant crosshead displacement rate of 2 mm*s^-1^. Interfacial adhesion strength was reported by dividing the maximum force by the initial area of contact between the PHB sheet and bio-sticker specimen. The energy dissipated was calculated by integrating the area under the force-displacement curve.

## Acknowledgments

The authors wish to thank Dr. Alyson Santoro, Dr. Melissa Omand, Marimikel Charrier, and Dr. Allison Pieja for advice and discussion about the project. Funding to A.S.M. was provided by the National Science Foundation via ITE-2137561 and ITE-2230641, by the Department of Energy via DE-SC0023354, and by the Arnold & Mabel Beckman Foundation. H.C. was supported by the U.S. Department of Energy, Office of Science, Basic Energy Sciences, under Award # DE-SC0019141. The scanning electron microscopy work was performed, in part, at the University of Rochester Integrated Nanosystems Center. Rheology testing was conducted at the University of Rochester’s Biomedical Engineering Department, Materials Mechanics Laboratory.

## Supporting Information

### Supporting Methods

*Logistic fitting of the viability and PHB degradation rates of 3D-bioprinted Bacillus sp. NRRL B-14911*.

The survival of 3D-bioprinted *Bacillus* sp. NRRL B-14911 was analyzed quantitatively. CFU data over time was fitted to a logistic bacterial growth equation^[32]^ (Equation 1) using the cftool app in MATLAB R2022b (MathWorks), to obtain estimates of growth rate (r), maximum growth rate (k), and initial growth rate (N).

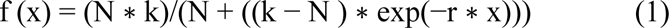

Parameter estimates with 95% confidence intervals are: N = 0.0002769 (0.0001325, 0.0004213), k = 0.05186 (0.04125, 0.06248), and r = 0.2659 (0.2283, 0.3034). The sum of squared estimate of errors (SSE) is 2.504e^−6^, R^2^ is 0.9981, and root mean square error is 0.0005595.

The PHB degradation rate of 3D-bioprinted bio-stickers containing *Bacillus* sp. NRRL B-14911 was analyzed quantitatively. PHB degradation data by bio-stickers over time with varying concentrations of PHB was fitted to a logistic growth equation (Equation 1) to obtain estimates of degradation rate (r), maximum degradation rate (k), and initial degradation rate (N). Parameter estimates with 95% confidence intervals are: N = 0.2002 (-1.607, 2.007), k = 6.685 (-1.092, 14.46), and r = 1.31 (-2.635, 5.255). SSE is 8.868, R^2^ is 0.6299, and root mean square error is 1.719.

**Supplemental Figure 1.**
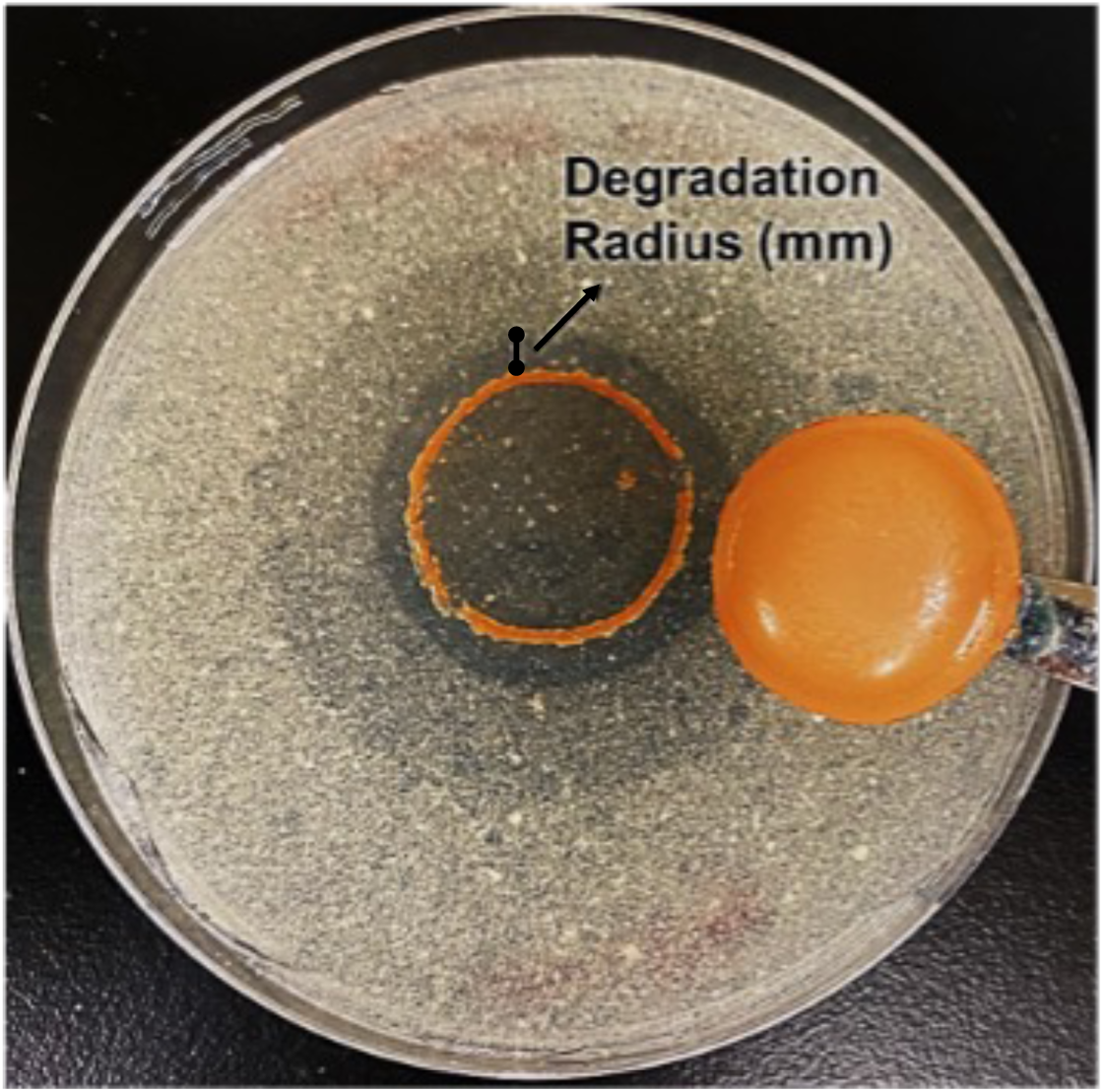
Clear-zone assay for PHB biodegradation. Degradation of PHB in a Marine Broth-agar plate by a bio-sticker (orange) containing 4% alginate and *Bacillus* sp. NRRL B-14911. PHB degradation can be visualized via the disappearance of opaque PHB powder and the appearance of clear zones within the agar.

**Supplemental Figure 2.**
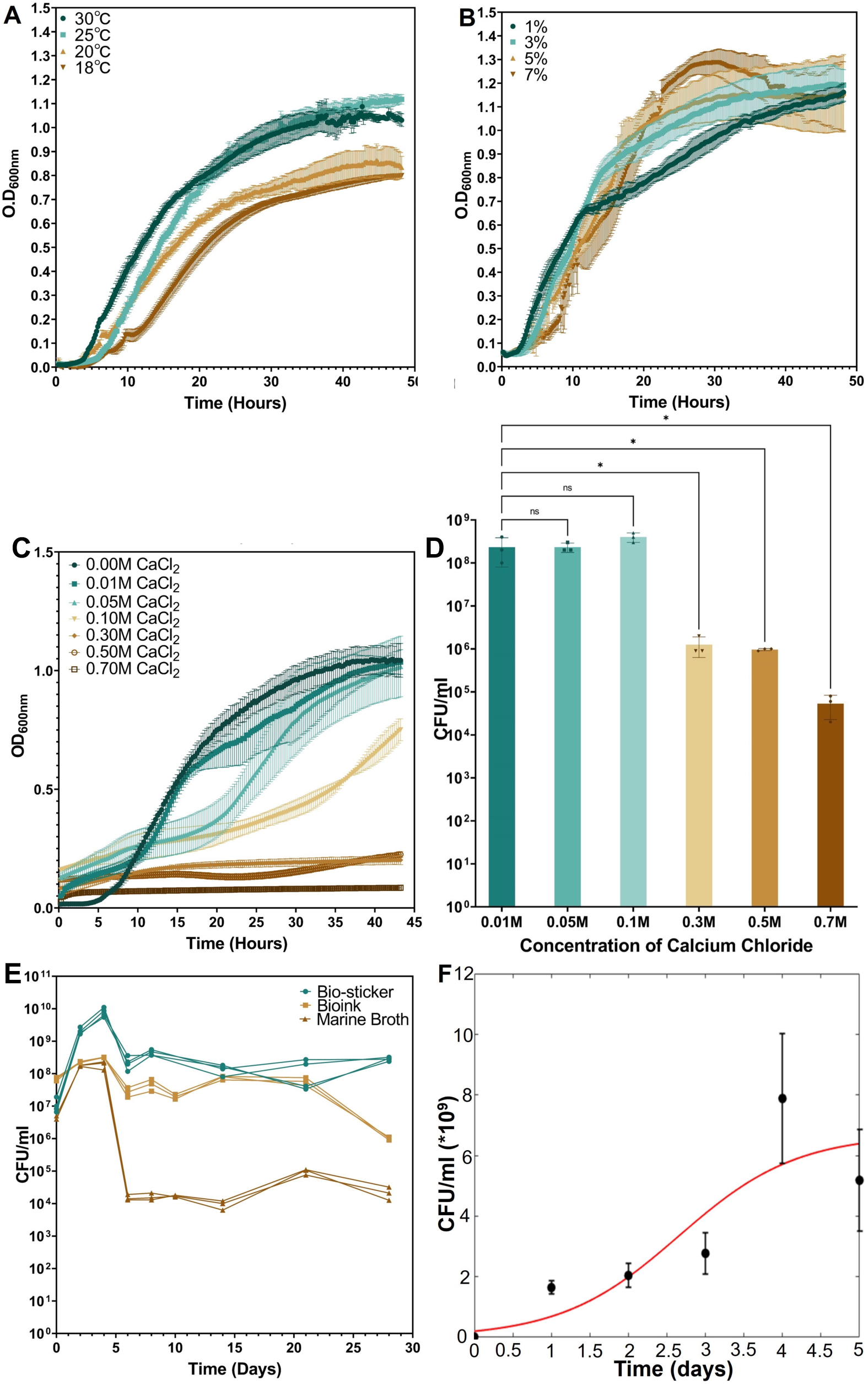
Growth and viability of *Bacillus* sp. NRRL B-14911 under different environmental and bio-printing conditions. (A) Growth curves of *Bacillus* sp. NRRL B-14911 cultured in Marine Broth at 30 °C (dark green), 25 °C (light green), 20 °C (light brown), and 18 °C (dark brown). (n=3) (B) Growth curves of *Bacillus* sp. NRRL B-14911 cultured in LB media supplemented with sodium chloride at 1% (dark green), 3% (light green), 5% (light brown), or 7% (dark brown). (n=3) (C) Growth curves of *Bacillus* sp. NRRL B-14911 cultured in Marine Broth supplemented with CaCl_2_ at concentrations ranging from 0.00 M (darkest green) to 0.70 M (darkest brown). (n=3) (D) CFUs of *Bacillus* sp. NRRL B-14911 in bio-stickers 3D-bioprinted onto Marine Broth-agar plates containing calcium chloride at concentrations ranging from 0.01 M (darkest green) to 0.7 M (darkest brown). (n=3) * P≤0.05, ns = not significant by one-way ANOVA statistical analysis. (E) CFU/mL of *Bacillus* sp. NRRL B-14911 following incubation in Marine Broth (dark brown), non-polymerized bio-ink (light brown), and 3D-bioprinted bio-stickers (dark green) over 28 days. (n=3). (F) CFU/mL of *Bacillus* sp. NRRL B-14911 over 5 days of growth within a 3D-biopriinted bio-sticker (black circles) was fit to a logistic equation. (n=3)

**Supplemental Figure 3.**
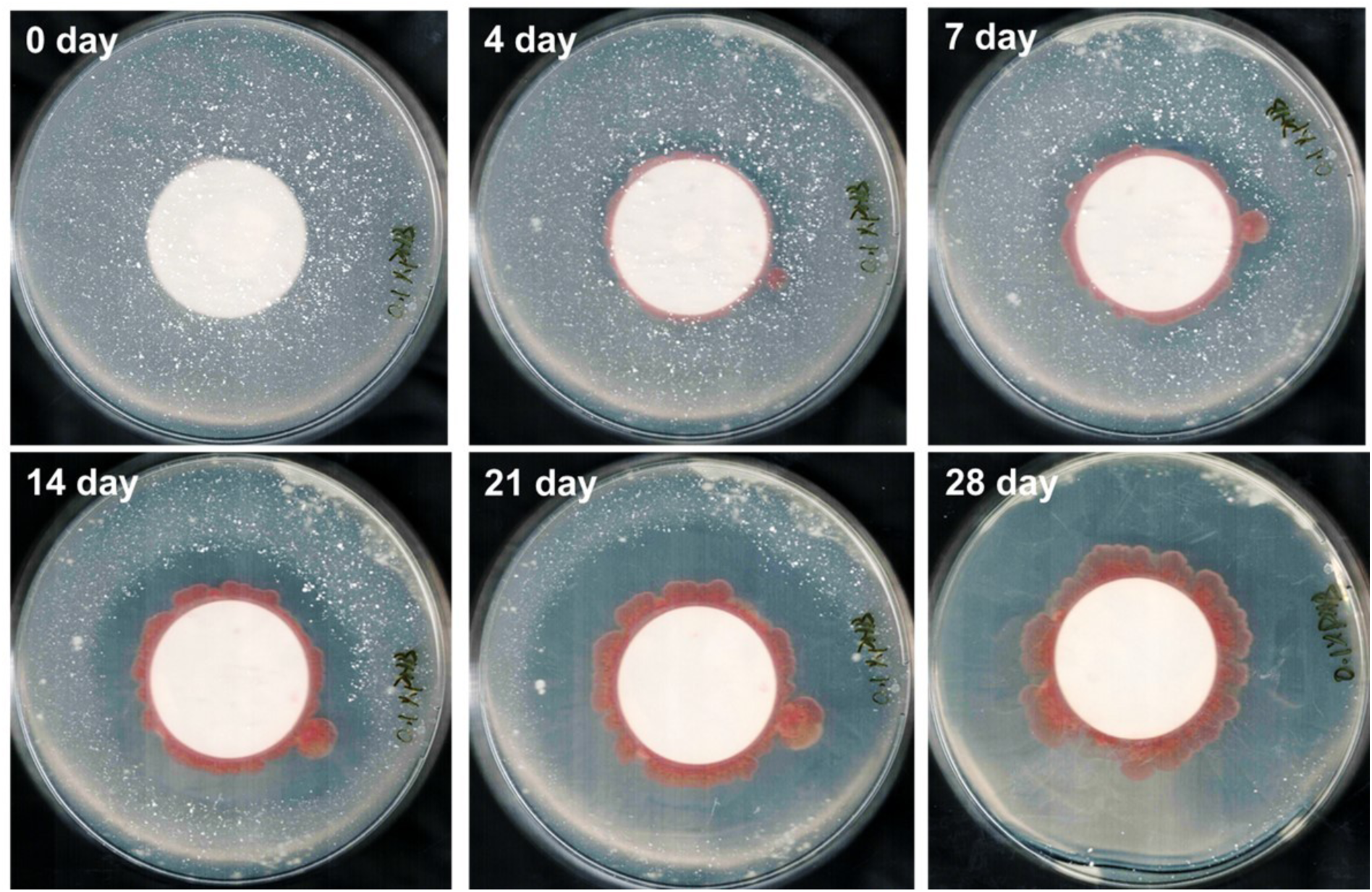
Preparation of samples for SEM analysis of bio-degradation of PHB sheets. Solid PHB discs (1-mm thickness, 44.5-mm diameter) were placed onto agar plates containing 0.3 M CaCl_2_ and 0.1% PHB powder. Bio-stickers containing *Bacillus* sp. NRRL B-14911 were applied overtop of the PHB discs, and samples were incubated at 30 °C for 28 days. At the end of the incubation period, the PHB discs were carefully removed, and residual bacteria and excess hydrogel were meticulously cleansed from their surfaces.

**Supplemental Figure 4.**
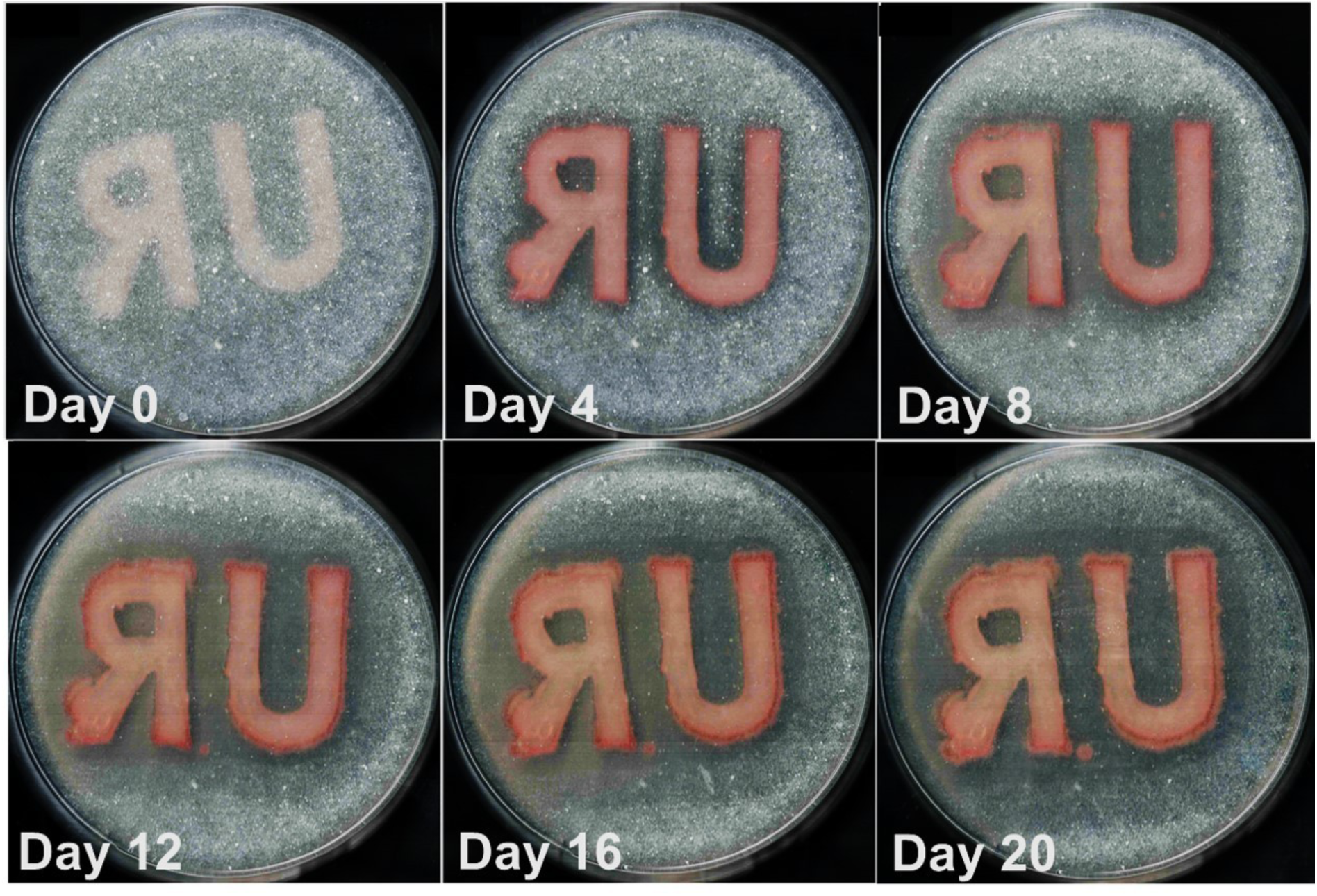
Progression of PHB degradation by bio-stickers with varied geometries. Bio-stickers containing *Bacillus* sp. NRRL B-14911 were 3D-printed into “U” and “R” shapes and placed onto Marine Broth-agar plates containing 0.3 M CaCl_2_ and 0.5% PHB powder. Samples were incubated at 30 °C for 28 days. During this time period, the PHB was first cleared from the agar underneath the bio-stickers, after which the clear zones continually expanded in radius.

**Supplemental Figure 5.**
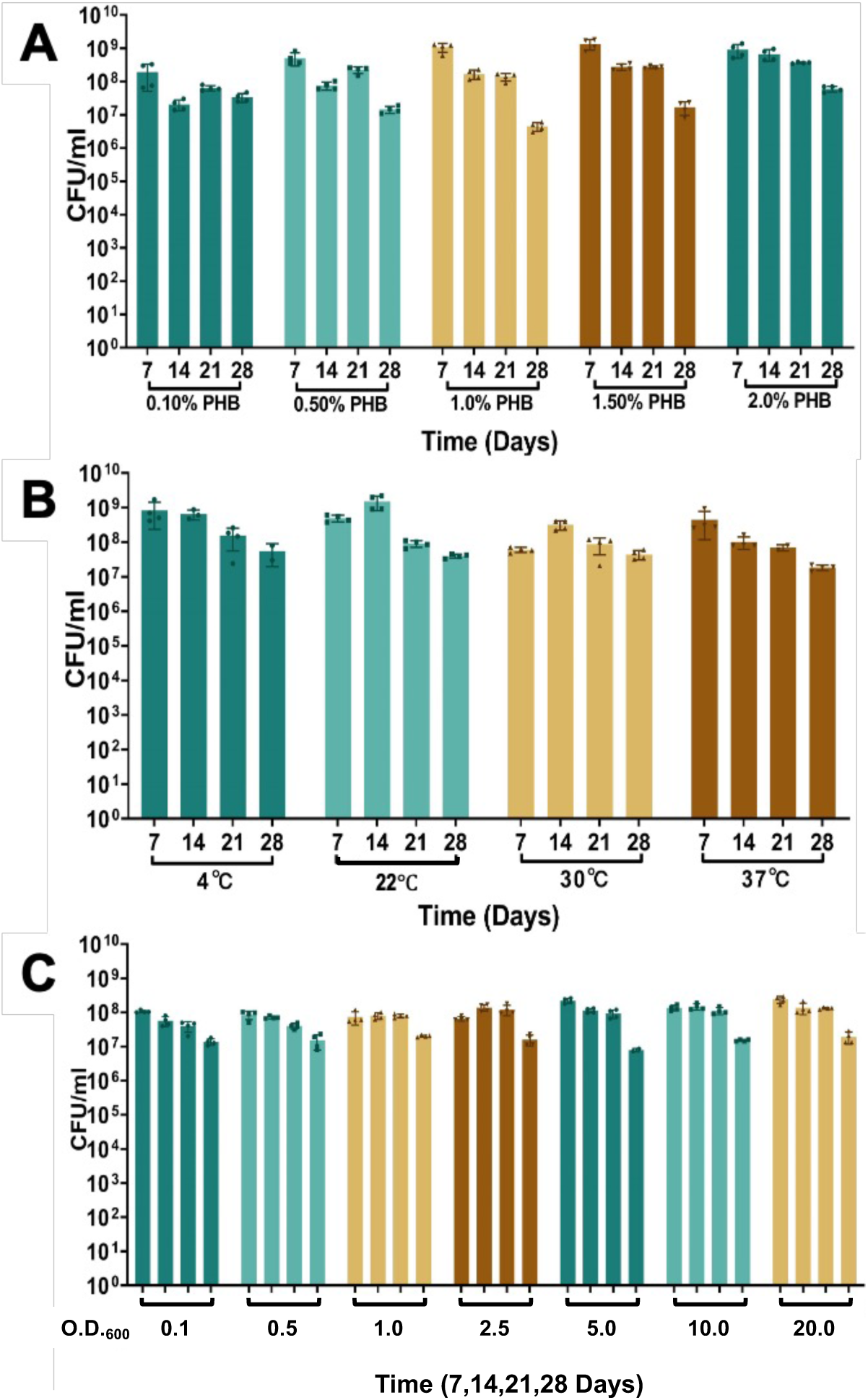
CFU values for bio-stickers with altered parameters. CFU assays for *Bacillus* sp. NRRL B-14911 3D-printed at an O.D._600_ of 10 into 10-mm diameter circular shapes and placed onto Marine Broth-agar plates containing 0.3 M CaCl_2_ and 0.1% PHB powder, followed by incubation at 30 °C, except where altered parameters are noted. (A) CFU/mL over 28 days for bio-stickers bio-printed onto plates containing varying PHB concentrations. (n=4) (B) CFU/mL over 28 days for bio-stickers incubated at varying temperatures. (n=4) (C) CFU/mL over 28 days for bio-stickers printed using bio-ink with varying initial O.D._600_. (n=4)

**Supplemental Figure 6.**
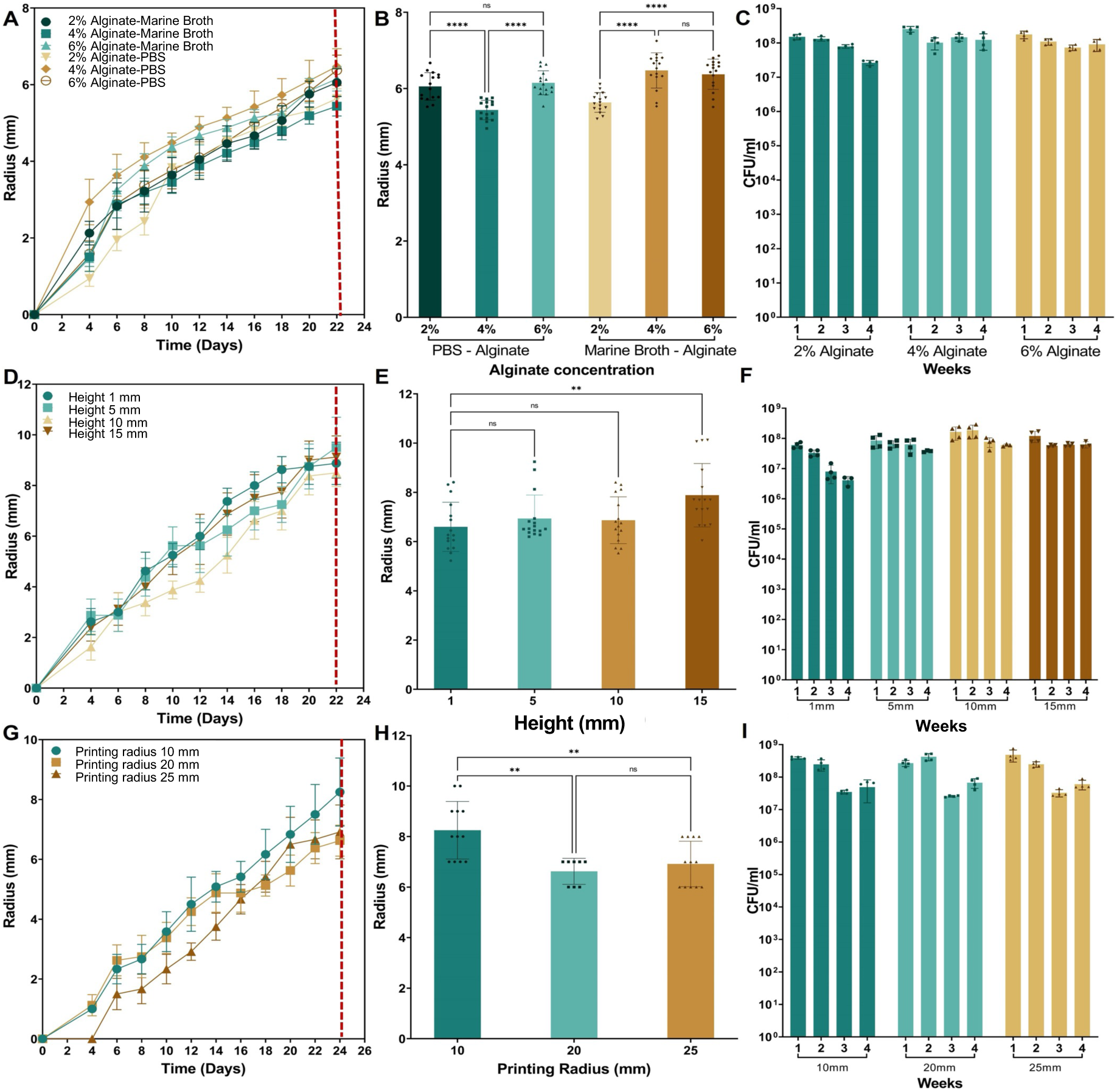
PHB degradation is modestly affected by tuning bio-sticker geometry and hydrogel density. Clear zone and CFU assays for *Bacillus* sp. NRRL B-14911 3D-printed at an O.D._600_ of 10 into 10-mm diameter circular shapes and placed onto Marine Broth-agar plates containing 0.3 M CaCl_2_ and 0.1% PHB powder, followed by incubation at 30 °C, except where altered parameters are noted. (A-C) Clear zone radius over time (A) and on day 22 (B) and CFU/mL over time (C) for bio-stickers containing varying alginate concentrations. CFU/mL data in panel C is for alginate-Marine Broth bio-stickers. (D-F) Clear zone radius over time (D) and on day 22 (E) and CFU/mL over time (F) for bio-stickers 3D-bioprinted with varying heights. (G-H) Clear zone radius over time (G) and on day 24 (H) and CFU/mL over time (I) for bio-stickers 3D-bioprinted with varying initial radii. Vertical dashed red lines in panels A, D, and G indicate time points that were analyzed in panels B, E, and H for statistical differences between conditions. ** P≤0.01, **** P≤0.0001, ns = not significant by one-way ANOVA statistical analysis

**Supplemental Figure 7.**
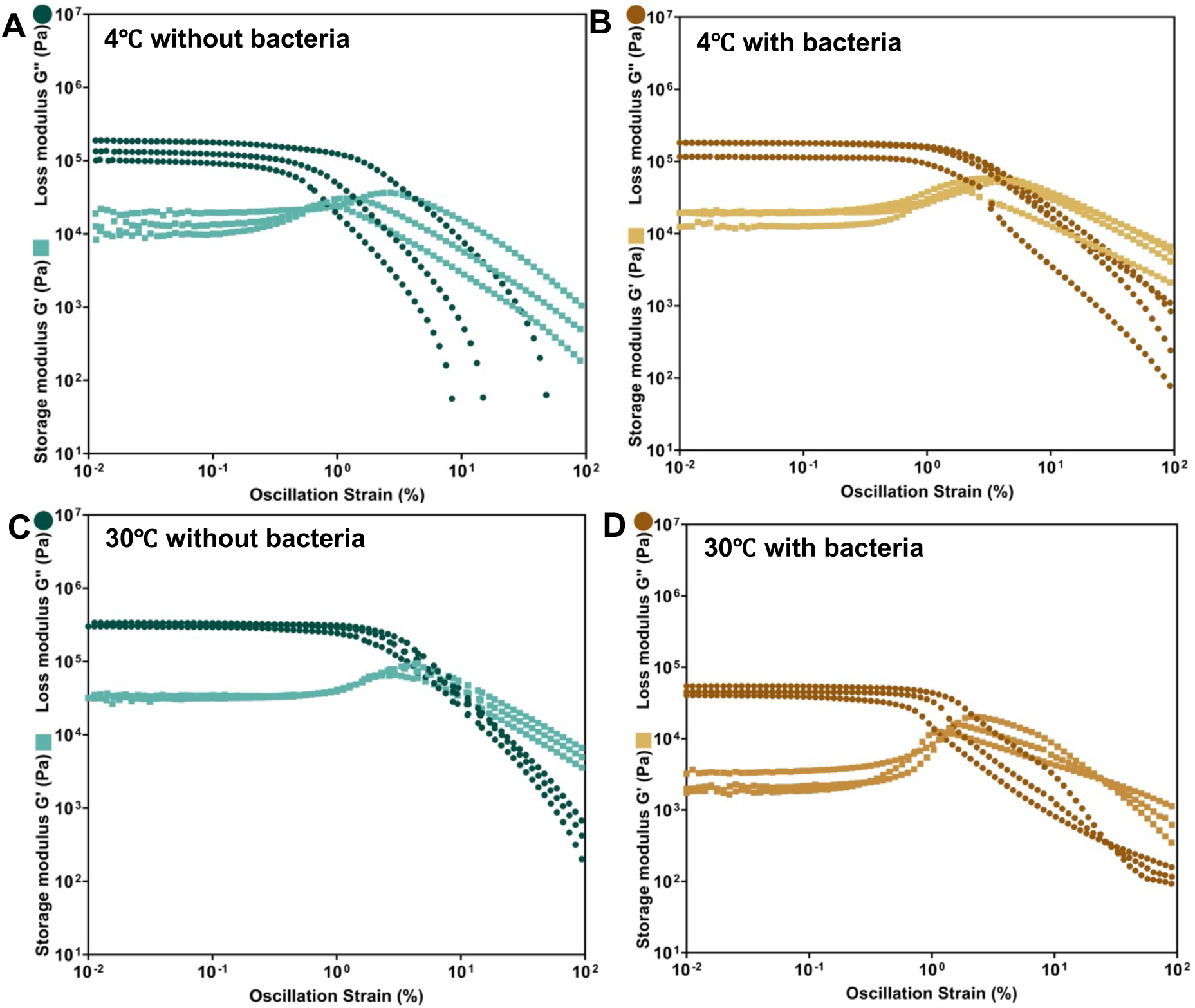
Rheometry data for bio-stickers. Rheometry testing data showing the relationship between oscillation strain and both storage modulus (squares) and loss modulus (circles) for (A-B) bio-stickers incubated at 4 °C and (A) not containing or (B) containing *Bacillus* sp. NRRL B-14911, and (C-D) bio-stickers incubated at 30 °C and (C) not containing or (D) containing *Bacillus* sp. NRRL B-14911.

**Supplemental Figure 8.**
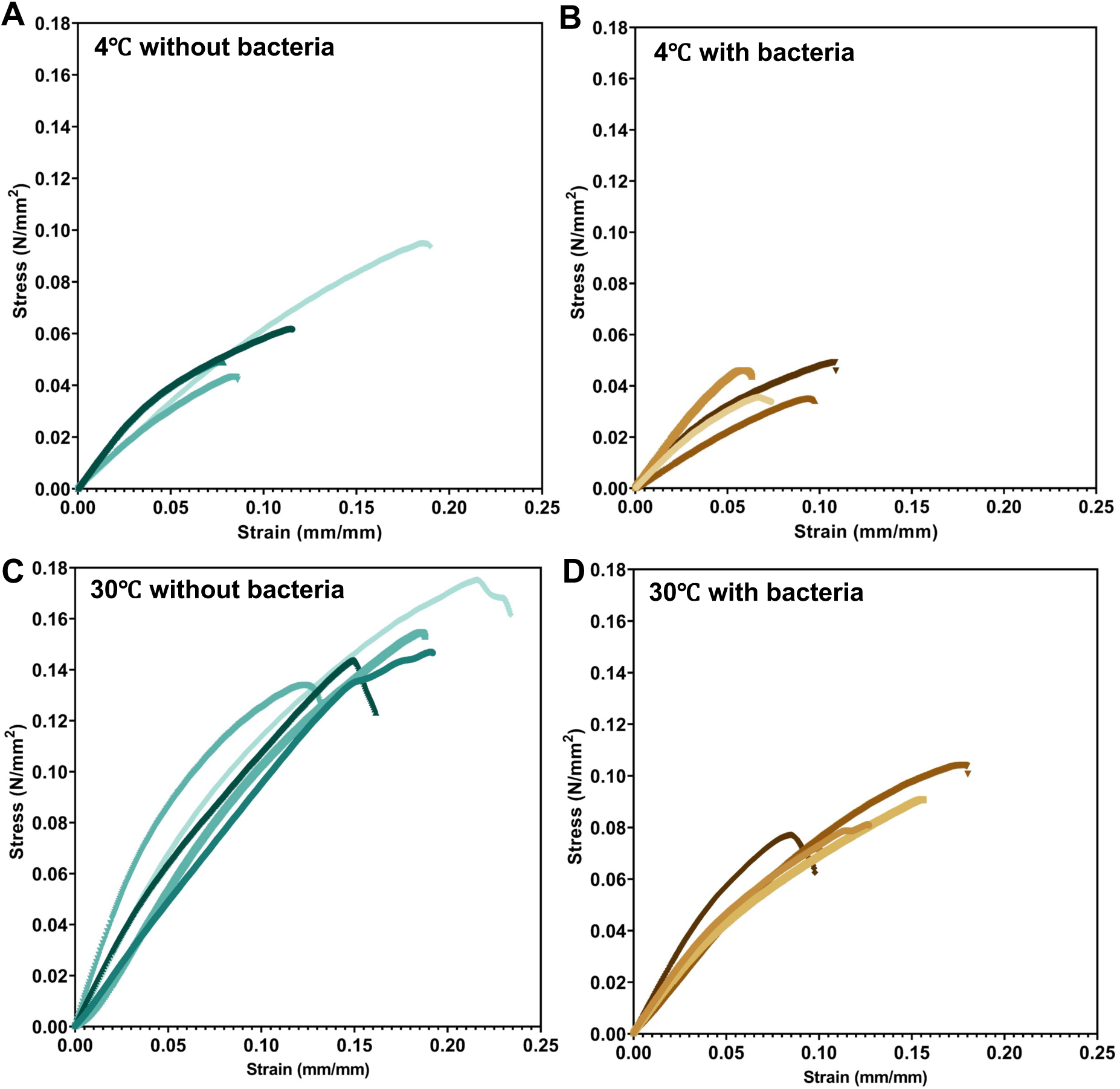
Tensile testing stress-strain curves for bio-stickers. (A-B) Tensile testing stress-strain curves for bio-stickers incubated at 4 °C and (A) not containing or (B) containing *Bacillus* sp. NRRL B-14911. (C-D) Tensile testing stress-strain curves for bio-stickers incubated at 30 °C and (C) not containing or (D) containing *Bacillus* sp. NRRL B-14911.

**Supplemental Figure 9.**
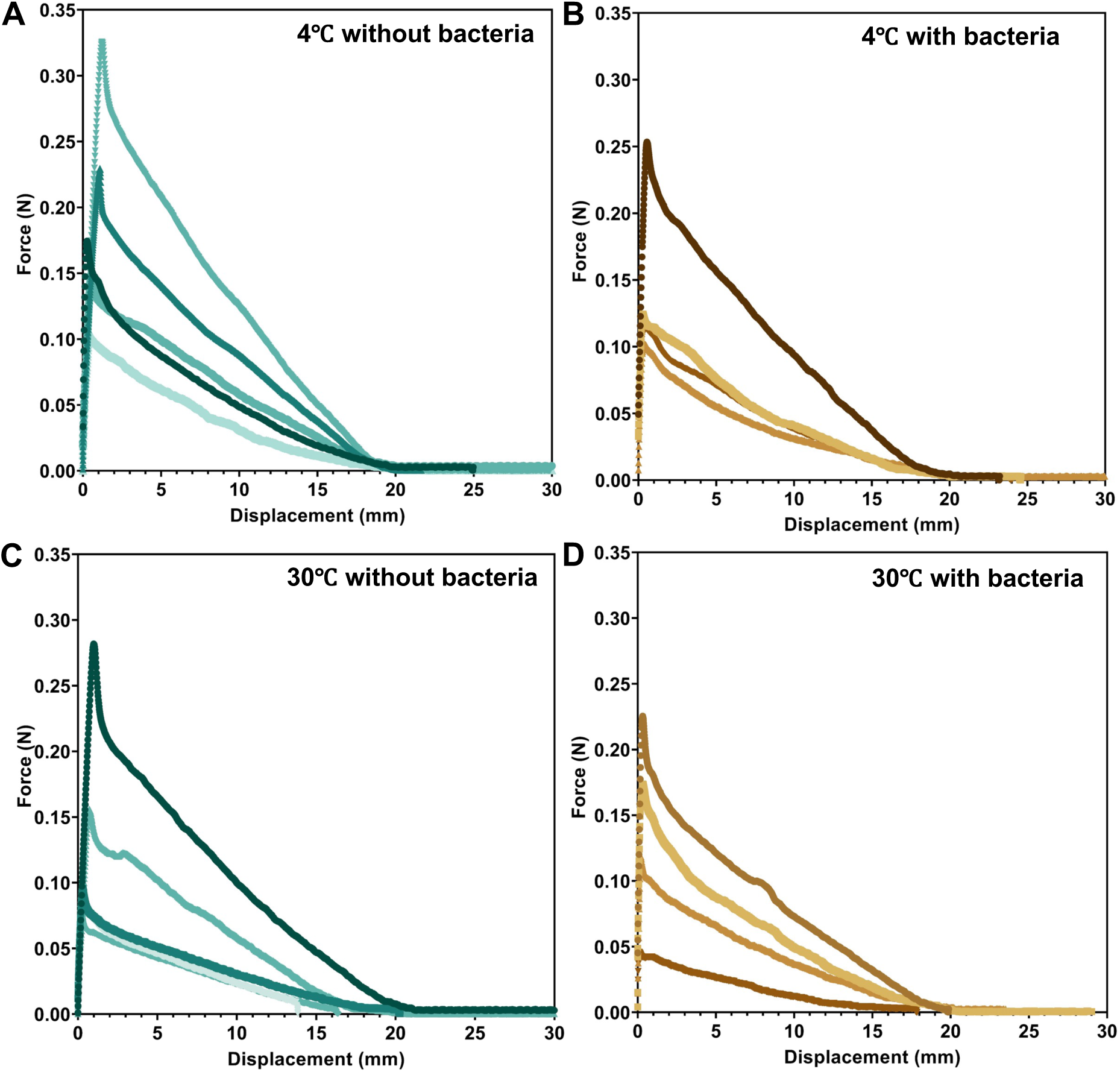
Adhesion testing force-displacement curves for bio-stickers. (A-B) Adhesion testing force-displacement curves for bio-stickers incubated at 4 °C and (A) not containing or (B) containing *Bacillus* sp. NRRL B-14911. (C-D) Adhesion testing force-displacement curves for bio-stickers incubated at 30 °C and (C) not containing or (D) containing *Bacillus* sp. NRRL B-14911.

## Notes

### Competing Interest Statement

A.S. Meyer is co-founder and holds equity in Nereid Biomaterials.

